# Optogenetic stimulation of medial amygdala GABA neurons with kinetically different channelrhodopsin variants yield opposite behavioral outcomes

**DOI:** 10.1101/2021.06.30.450543

**Authors:** Aiste Baleisyte, Ralf Schneggenburger, Olexiy Kochubey

## Abstract

Optogenetic manipulation of genetically-specified neuron populations has become a key tool in circuit neuroscience. The medial amygdala (MeA) receives pheromone information about conspecifics and has crucial functions in social behaviors; interestingly, this amygdalar structure contains a majority of GABAergic projection neurons. A previous study showed that optogenetic activation of MeA GABA neurons with channelrhodopsin-2^H134R^ (ChR2) strongly enhanced inter-male aggression (Hong et al. 2014, Cell). When we attempted to reproduce these findings, accidentally using a faster channelrhodopsin variant (channelrhodopsin-2^H134R,E123T^ or ChETA), we found the opposite results. We therefore systematically compared the behavioral outcome of optogenetic stimulation of MeApd GABA neurons with ChETA versus ChR2, employing two widely used AAV serotypes. This revealed that optogenetic stimulation with ChETA *suppressed* aggression, whereas optogenetic stimulation with ChR2 *increased* aggression. Recordings of membrane potential changes following optogenetic stimulation with ChETA versus ChR2 revealed larger plateau depolarizations, smaller action potential amplitudes, and larger local inhibition of neighboring inhibitory neurons with ChR2 as compared to ChETA. Our study shows that channelrhodopsin variants have to be chosen with care for *in-vivo* optogenetic experiments. Furthermore, the role of MeApd GABA neurons in aggression control should be re- evaluated.

## Introduction

The advent of optogenetic methods about a decade ago has allowed researchers to manipulate the electrical excitability and AP-firing of genetically-specified neuron populations by light, and to observe the influence of this manipulation on animal behavior (Deisseroth, 2015; Tye & Deisseroth, 2012; Zhang et al., 2007). The “excitatory” cation channel channelrhodopsin-2 from *Chlamydomonas reinhardtii* has been amongst the first light-sensitive ion transporters from single-cell organisms made available for optogenetic studies (Boyden et al., 2005; Nagel et al., 2002, 2005; Sineshchekov et al., 2002). After its initial description, the properties of channelrhodopsin-2 were optimized for *in-vivo* use by introducing engineered mutations. A histidine to arginine mutation was introduced to increase the stationary photocurrent (H134R; Nagel et al., 2005), giving rise to the channelrhodospin-2^H134R^ variant (called here “ChR2”), which has been widely used for *in-vivo* optogenetic stimulation experiments. In subsequent work, additional mutations were introduced with the aim to increase the off-kinetics of the light-gated channel; one of the resulting variants carries an additional E123T mutation (channelrhodopsin-2^H134R, E123T^) and has been called “ChETA” (Gunaydin et al., 2010). In addition, other light-gated ion channels with fast kinetics have become available, like for example Chronos and ChIEF (Klapoetke et al., 2014; Lin et al., 2009). The kinetic differences between ChR2, and the faster ChETA, ChIEF and Chronos have been well documented (Berndt et al., 2011; Gunaydin et al., 2010; Klapoetke et al., 2014; Lin et al., 2009; Mattis et al., 2012), and it was shown that blue light stimulation of ChR2 expressing neurons can lead to a light-induced depolarization block, especially in some classes of interneurons (Herman et al., 2014). Nevertheless, fundamentally conflicting behavioral outcomes after the *in-vivo* use of different channelrhodopsins have, to our knowledge, not been found.

The medial amygdala, and especially its postero-dorsal division (MeApd) contains a large fraction of GABA neurons (Choi et al., 2005; Keshavarzi et al., 2014). Of these, many neurons are long-range projection neurons that send axons to hypothalamic nuclei and other targets (Bian et al., 2008; Canteras et al., 1995; Choi et al., 2005; Keshavarzi et al., 2014). Early lesion studies showed a role of the MeA in social behavior and aggression control (Miczek et al., 1974; Vochteloo and Koolhaas, 1987; reviewed in Haller, 2018). More recent optogenetic and chemogenetic studies in mice have further defined the role of the MeA in aggression control (Hong et al., 2014; Nordman & Li, 2020; Padilla et al., 2016; Unger et al., 2015). Particularly, Hong et al. (2014) have shown, using a *VGAT^Cre^* mouse line to target GABA neurons in the MeApd, that optogenetic stimulation of these neurons using ChR2 led to a marked increase of attacks in the resident-intruder test (Hong et al., 2014). This finding has given rise to the generally accepted view that MeApd GABA neurons have a *stimulatory* role in aggression control (see reviews by Aleyasin et al., 2018; P. Chen & Hong, 2018; Hashikawa et al., 2016; Lischinsky & Lin, 2020).

We now report that using ChR2 and ChETA for optogenetic stimulation of MeApd-GABA neurons gives rise to a paradoxical *opposite* behavioral outcome in a social behavior test. In initial experiments in which we wished to follow-up on the role of MeApd GABA neurons in aggression control, we accidentally used ChETA and observed an *inhibition* of aggression, opposite to the previous study that had employed ChR2 (Hong et al., 2014). We therefore systematically compared the behavioral effects of optogenetic stimulation with ChR2 *versus* ChETA. This revealed that under otherwise identical conditions, activating MeApd GABA neurons with ChETA *suppressed* aggression, whereas optogenetic stimulation with ChR2 led to an *increase* of aggression as reported earlier (Hong et al., 2014). *Ex-vivo* electrophysiological recordings showed that optogenetic stimulation of MeApd GABA neurons expressing the slower ChR2 showed significantly larger plateau depolarizations than with ChETA. The plateau depolarizations caused a time-dependent decrease of AP amplitudes during prolonged stimulation trains, and a larger charge transfer during optogenetically-evoked inhibition of neighboring MeApd GABA neurons, as compared to stimulation using ChETA, differences which might explain the opposite behavioral outcomes *in-vivo*. Our study shows that care has to be taken when selecting channelrhodopsin variants for *in-vivo* optogenetic experiments, especially in mixed populations of GABA neurons. Given these conflicting results with two channelrhodopsin variants, the role of MeApd GABA neurons in controlling aggressive behavior should be newly investigated.

## Results

### Optogenetic stimulation of MeApd GABA neurons with ChETA reduces aggression

Following a previous pioneering study which showed a *stimulatory* role of MeApd GABA neurons in the control of aggression (Hong et al., 2014), we originally wished to investigate the role of molecularly - and projection-defined MeApd GABA neurons in this social behavior. For this, we first attempted to reproduce the earlier findings, in which MeApd GABA neurons were optogenetically stimulated using the VGAT^Cre^ mice, leading to targeting the entire GABA neuron population (Figures 1A, B; see also Hong et al., 2014). Male *VGAT^Cre^ x tdTomato* mice were bilaterally injected with an AAV vector driving the Cre-dependent expression of ChETA (AAV8:hSyn:DIO:ChETA-eYFP; see Table 1). Control mice received a vector driving the expression of eGFP (AAV8:hSyn:DIO:eGFP), but underwent otherwise identical procedures. Optic fibers were implanted bilaterally in the MeA, targeting the postero-dorsal subnucleus (Figure 1B; Figure 2). Four weeks later, the mice were subjected to a resident-intruder test in their home cage using an adult male BALB/cByJ mouse as an intruder. We used trains of blue light pulses (5 ms, 10 mW, 20 Hz, length of 30s or 60s) delivered at regular intervals (60s dark periods), to avoid a possible bias by experimenter-driven stimulation.

**Figure 1.**
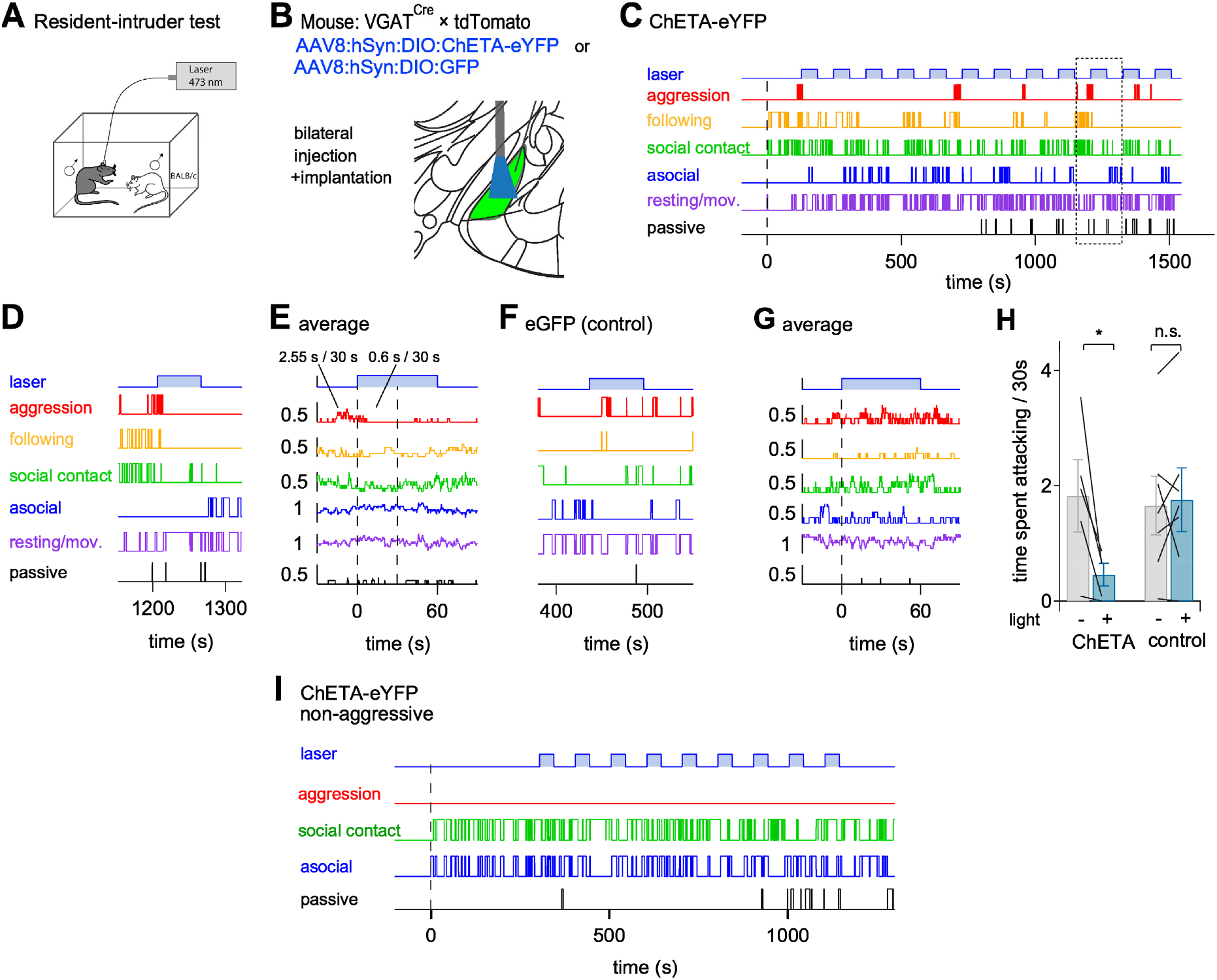
In-vivo optogenetic stimulation of MeApd GABA neurons using ChETA inhibits inter-male aggression. (A) Scheme of a resident-intruder test with optogenetic stimulation of the MeApd in the resident male. BALB/cByJ male mice (one at a time) were used as intruders. (B) Schematic of the experimental approach for expression of ChETA and optic fiber implantation targeting. (C) Traces showing various quantified behaviors of a resident mouse during a resident-intruder test (see Materials and Methods). An intruder was introduced at t = 0. Trains of laser light pulses (5 ms, 20 Hz repetition rate, 10 mW intensity) were applied for 60s, interleaved by 60s dark periods. Note that the aggression bouts (red trace) preferably occurred *outside* the light trains. (D) An example light train episode marked by the dotted rectangle in (C). (E) Aligning the behavior traces to the onsets of light trains with subsequent averaging showed a reduction of aggression after the start of the light train. The average time spent attacking during the last 30s of darkness, and during the first 30s of light stimulation, is indicated. Data in (C-E) are from a single ChETA-expressing example mouse. (F, G) Behavioral data from a control mouse expressing eGFP, both before and during a single light train (F) and corresponding average behavior traces for n = 7 light trains (G). As expected, in this eGFP-expressing control mouse, blue-light stimulation did not modulate aggressive behavior. (H) Quantification of the time spent attacking during the 30 s before (-), and 30 s into (+) the light stimulation in mice expressing ChETA (N = 5) or eGFP (N = 7). For each mouse, the individual values are averages across two tests performed on subsequent days. Note the significant inhibition of aggression by blue light in the ChETA group (p = 0.0385), but not in the eGFP group (p = 0.73). (I) An example mouse expressing ChETA, which showed no aggression against the intruder (non-aggressive mice were not included in in the data pool shown in H). Note that optogenetic stimulation with ChETA did not induce aggression. Similar observations were made in N = 3 non-aggressive mice expressing ChETA. Error bars are ± s.e.m. See also Suppl. Figure 1.

**Figure 2.**
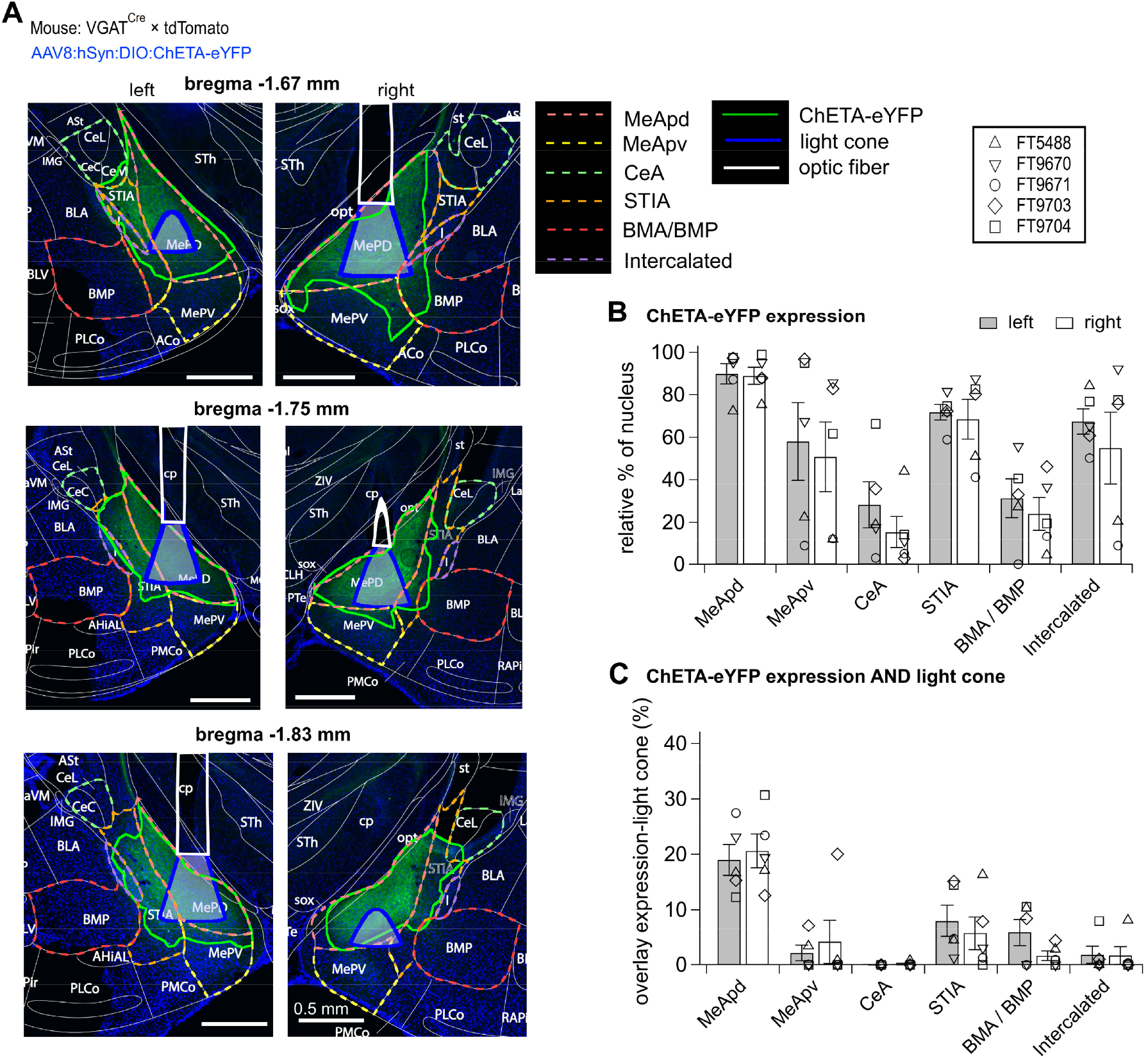
Histological reconstructions of ChETA-eYFP expressing brain areas and optic fiber placement confirm optogenetic stimulation of the MeApd. (A) Fluorescent images on the level of the left and the right MeApd (bregma levels indicated), obtained from a ChETA-expressing mice after the behavior experiments (Figure 1). The regions of ChETA-eYFP expression (fluorescence in green channel) were outlined (solid green lines). Reconstructed contours of cross-sections through the optic fiber (solid white lines) and the brain atlas outlines (dashed lines; Franklin and Paxinos, 2013) were overlaid. The area of effective illumination (blue shading) was derived as explained in Materials and Methods. (B) The percentage of the area expressing ChETA-eYFP in the MeApd and neighboring brain structures, integrated across serial brain sections (N = 5 mice). See Results for abbreviations. (C) The percentage of each nucleus which expressed ChETA-eYFP *and* overlapped with the effective illumination area (see A). Note that, despite an expression of ChETA-eYFP expression in some neighboring structures (B), the volume of efficient optogenetic stimulation is predominantly located in the MeApd.

**Table 1.**
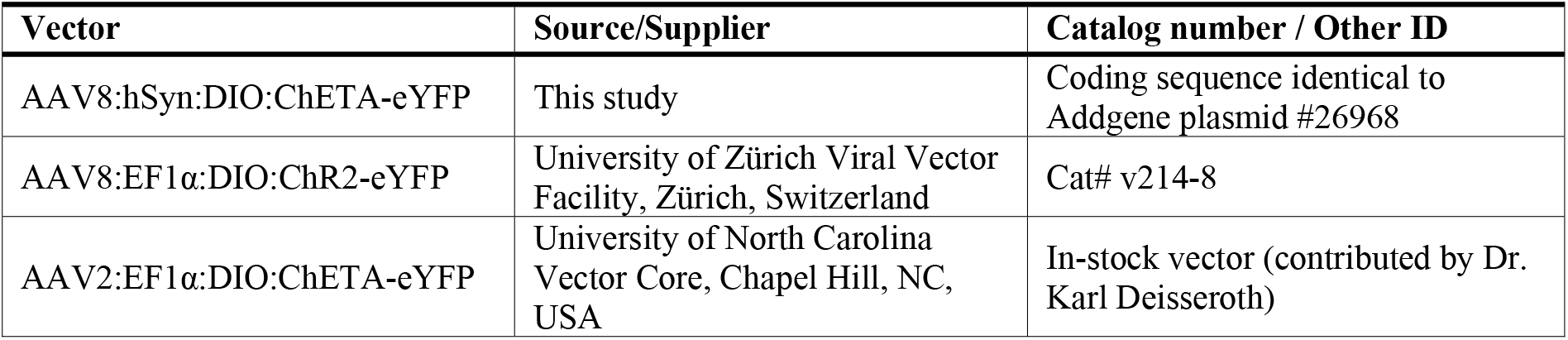
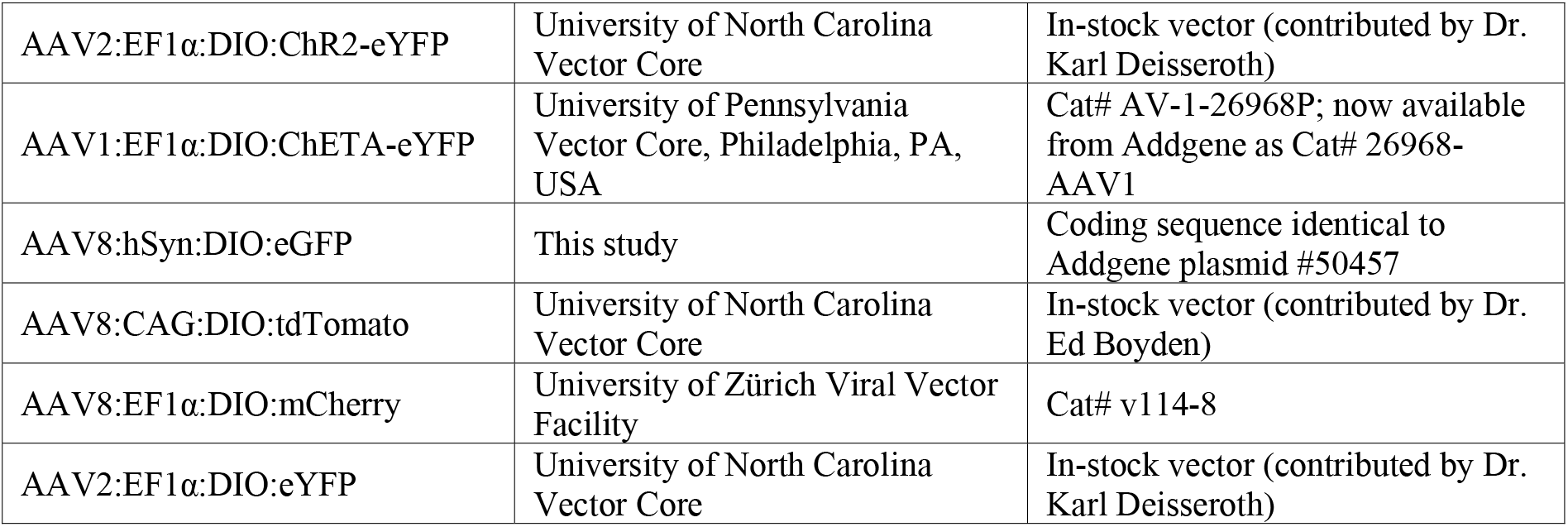
List of the viral vectors used in the study.

| Vector | Source/Supplier | Catalog number / Other ID |
| --- | --- | --- |
| AAV8:hSyn:DIO:ChETA-eYFP | This study | Coding sequence identical to Addgene plasmid #26968 |
| AAV8:EF1α:DIO:ChR2-eYFP | University of Zürich Viral Vector Facility, Zürich, Switzerland | Cat# v214-8 |
| AAV2:EF1α:DIO:ChETA-eYFP | University of North Carolina Vector Core, Chapel Hill, NC, USA | In-stock vector (contributed by Dr. Karl Deisseroth) |
| AAV2:EF1α:DIO:ChR2-eYFP | University of North Carolina Vector Core | In-stock vector (contributed by Dr. Karl Deisseroth) |
| AAV1:EF1α:DIO:ChETA-eYFP | University of Pennsylvania Vector Core, Philadelphia, PA, USA | Cat# AV-1-26968P; now available from Addgene as Cat# 26968-AAV1 |
| AAV8:hSyn:DIO:eGFP | This study | Coding sequence identical to Addgene plasmid #50457 |
| AAV8:CAG:DIO:tdTomato | University of North Carolina Vector Core | In-stock vector (contributed by Dr. Ed Boyden) |
| AAV8:EF1α:DIO:mCherry | University of Zürich Viral Vector Facility | Cat# v114-8 |
| AAV2:EF1α:DIO:eYFP | University of North Carolina Vector Core | In-stock vector (contributed by Dr. Karl Deisseroth) |

We observed that aggression bouts occurred mostly in the dark periods (60s) in-between light stimulation, and that blue light often led to a stop of aggression (Figures 1C-1E). Aligning and averaging the behavioral scores from individual mice showed that blue light reduced the time that a mouse spent attacking, from 2.55 s per 30 s of dark period (averaged over n = 10 dark periods), to 0.6 s per 30s of light period (n = 10; Figure 1E). Blue light stimulation reduced the time mice spent attacking in all animals of the ChETA group (Figure 1H; p = 0.039, N = 5 mice; two-tailed paired t-test). Control mice expressing eGFP did not show a modulation of aggression by light, as shown in Figures 1F, G for an example animal (N = 7 mice, p = 0.73; two-tailed paired t-test, Figure 1H). These findings, unexpectedly, suggest that optogenetic stimulation of MeApd GABA neurons leads to a *decrease* of aggression, opposite to the previous results (Hong et al., 2014).

Our experimental mice were single-housed after surgery and were not further selected according to their aggression levels (see Materials and Methods). In some mice, we found low levels of aggression and no spontaneous attacks against the intruder. In none of these non-aggressive mice did optogenetic activation of MeApd GABA neurons induce attacks (see Figure 1I for an example; N = 3 non-aggressive mice). This finding is also different from the previous study, in which blue light readily triggered attacks in non-aggressive mice (Hong et al., 2014; their Figure 2). Furthermore, in an additional N= 2 mice in which ChETA was expressed by an AAV1 serotype vector, we similarly observed that blue light suppressed aggression (Suppl. Figure 1). In the latter experiments, we also varied the intensity of blue light stimulation, and observed that the inhibition of aggression depended on light intensity, with no other obvious behaviors stimulated at low light intensities (Suppl. Figure 1). These observations further support our finding that optogenetic stimulation of MeApd - GABA neurons *inhibits* aggression (Figure 1), opposite to the previous report (Hong et al., 2014).

A factor that may explain the opposite outcome of optogenetic stimulation between the present and the previous study (Hong et al., 2014) is that different brain areas might have been targeted between the two studies. We therefore reconstructed the regional expression of ChETA-eYFP, and the fiber placements for the experimental mice shown in Figures 1C- 1I (Figure 2A; see Materials and Methods). We found that 90 ± 5% and 89 ± 4 % of the cumulative area of the MeApd located in the vicinity of the optic fiber expressed ChETA (Figures 2A and 2B; for the left and the right brain side respectively, N = 5 mice). The expression of ChETA was not limited to the MeApd but extended into the adjacent areas of neighboring subnuclei of the amygdala, albeit filling smaller percentages of those areas (Figures 2A, 2B; MeApv, posterioventral MeA; CeA, central amygdala; STIA, bed nucleus of the stria terminalis, intraamygdaloid division; BMA/BMP, basomedial amygdaloid nuclei, anterior or posterior; I, intercalated nuclei of amygdala). Next, we estimated the percentages of the ChETA-positive brain areas which, based on the position and angle of the optic fiber, were likely illuminated by blue light (see Materials and Methods). The largest fractional overlap between the area of ChETA expression and the calculated light cone was found for the MeApd (19.0 ± 2.8 % and 20.7 ± 3.1 % for the left and right brain side; Figure 2C), whereas the overlap was negligible in the MeApv, CeA, and BMA (∼3.1%, ∼0.08% and ∼3.8%, respectively). The STIA region had a somewhat larger overlap of ChETA expression and illumination (8.0 ± 2.9 % and 5.7 ± 3.0 % of STIA volume for the left and right brain side; Figure 2C). However, it is possible that a significant part of ChETA-eGFP positive tissue in STIA originates from axons that project away from the MeApd. Taken together, post-hoc histological validation showed that we have primarily targeted the MeApd. Thus, we conclude that optogenetic stimulation of MeApd GABA neurons, using ChETA expressed by an AAV8 vector, suppresses aggression (Figures 1, 2).

### Optogenetic stimulation of MeApd GABA neurons with ChR2, independent of AAV serotype, leads to increased aggression

We next looked for other factors that might be different between our experiments (Figures 1, 2) and the previous study (Hong et al., 2014). The previous study had used an AAV2 serotype, driving the expression of the slower channelrhodopsin variant ChR2 (channelrhodopsin-2^H134R^; see Introduction). We therefore next performed optogenetic stimulation experiments of MeApd GABA neurons, using AAV2:EF1α:DIO:ChR2-eYFP (Figure 3A), a commercial vector as close as possible to the previously used custom construct (see Materials and Methods). With this vector, under otherwise identical conditions as in Figure 1, we found that optogenetic activation of MeApd GABA neurons *stimulated* attack behavior. In an example animal, the time spent attacking increased from an average of 0.22 s per 30 s of dark periods, to an average of 4.86 s per 30 s of light-stimulation period (Figures 3B-D). Across all mice in which we employed AAV2:ChR2, blue-light stimulation significantly increased aggression (Figure 3G, left; N = 7 mice; p = 0.012; two-tailed paired t-test). On the other hand, in control mice expressing eYFP under an AAV2 vector, stimulation with blue light did not modulate the attack frequency (Figure 3G, right; p = 0.35). One of the mice did not show spontaneous baseline aggression. In this mouse, in remarkable contrast to our results with ChETA (Figure 1I), the first 2-3 blue light stimulation trains triggered time-locked attacks (Figure 3H). Histological reconstructions showed that we had successfully targeted the MeApd in the experiments with AAV2:ChR2 (Suppl. Figure 2), quantitively similar to our previous experiments with ChETA (see above, Figure 2). Therefore, the experiments with AAV2:ChR2 in Figure 3 suggest that with ChR2 for optogenetic stimulation, the behavioral outcome is opposite from what is observed with ChETA.

**Figure 3.**
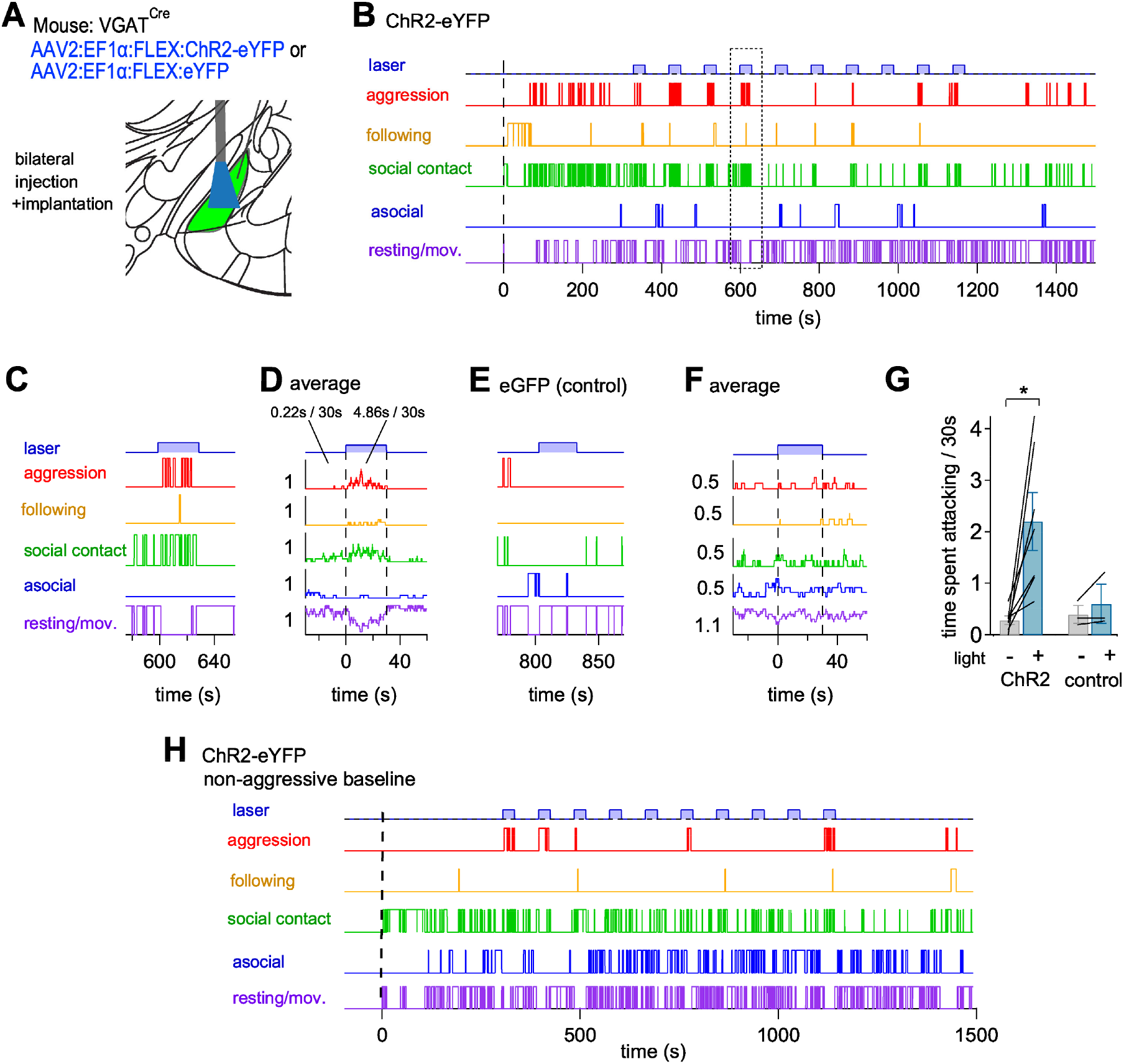
In-vivo optogenetic activation of MeApd GABA neurons using ChR2 stimulates inter-male aggression. (A) Scheme of the approach to express ChR2-eYFP and for optic fiber placement. (B) Behavior traces during the resident-intruder test with a mouse expressing ChR2 in the MeApd GABA neurons. 30s-long trains of blue-light pulses (1 ms, 20 Hz, 10 mW) were interleaved with 60 s dark periods. Note that under these conditions, attacks occurred preferentially *during* illumination, in contrast to the experiments with ChETA (Figure 1). (C - D) Example light episode from (B) marked with a dotted rectangle (C), and average behavior traces after alignment to the light onsets for n = 10 consecutive light trains (D). Numbers indicate average time spent attacking 30s before the light, and during the 30s light train. Note the increase of aggression during the light stimulation. (E-F) Behavior data for a VGAT^Cre^ mouse expressing eYFP (a control mouse). (G) Quantification of the effect of optogenetic stimulation on aggression in mice that expressed ChR2 (N=7) as compared to eYFP (N=3 control mice). Individual data are averages across two resident-intruder tests made on subsequent days. (H) An example mouse expressing ChR2, which was initially non-aggressive against the intruder mouse. Note that from the first trains of blue light pulses, this mouse started attacks against the intruder mouse. This observation further indicates that optogenetic activation of MeApd GABA neurons under ChR2 *stimulates* aggression. Error bars are ± s.e.m. See also Suppl. Figure 2.

The experiments performed in Figure 1 and 3, were, however, also different in terms of the AAV serotype. Thus, while we had performed our initial experiments with an AAV8 serotype (Figure 1), we used an AAV2 serotype in Figure 3 to allow for a direct comparison with the previous study (Hong et al., 2014). Additional experiments showed that AAV2 infected a larger number of MeApd GABA neurons than AAV8 (207 ± 19 %; n = 7 sections from N = 3 mice; Suppl. Figure 3). We found that only few neurons were infected by AAV8 *alone* and thus, AAV8-transduced neurons were essentially a subpopulation of the AAV2- transduced MeApd GABA neuron population (see Venn diagrams in Suppl. Figure 3).

To investigate whether the AAV serotype, or rather the expressed channelrhodopsin variant (ChR2 versus ChETA) determined the behavioral outcome, we next performed experiments with the two missing combinations of AAV serotypes and channelrhodopsin variants. These experiments unambiguously showed that the channelrhodopsin variant, but not the AAV serotype, determined the behavioral outcome (Suppl. Figure 4, and Figure 4). Thus, when ChETA was expressed with an AAV2 vector, a significant *decrease* in aggression was observed (Figure 4, third dataset; N = 9 mice; p = 0.0039; Wilcoxon two-tailed test; see Suppl. Figures 4A-4D for an example mouse). This finding thus confirms the results of Figure 1 with AAV8:ChETA; the latter results are re-plotted in Figure 4, leftmost dataset. Conversely, when we expressed ChR2 using an AAV8 vector, blue light led to a significant *increase* in aggression as compared to dark periods (Figure 4, fourth data set from left; N = 8 mice; p = 0.0391; Wilcoxon two-tailed test; see Suppl. Figures 4E-4G for an example mouse). This experiment confirms our results in Figure 3 with AAV2:ChR2; the latter results are re-plotted in Figure 4, second dataset. In mice expressing the inert eGFP or eYFP under the AAV8 or AAV2 serotype respectively, blue light stimulation was inefficient as expected (Figure 4, fifth and sixth datasets; N = 11 and 7 mice for AAV8 and AAV2 respectively; p = 0.41 and 0.53; two-tailed Wilcoxon test and paired t-test).

**Figure 4.**
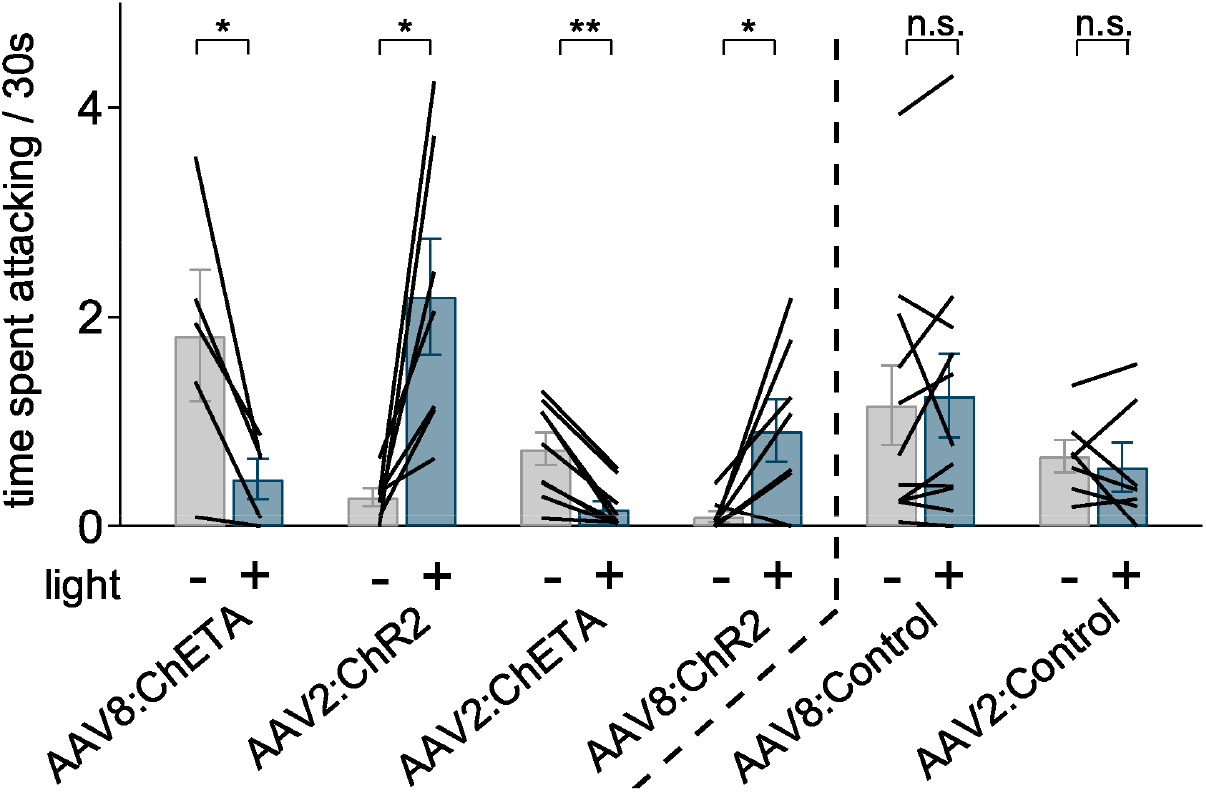
Channelrhodopsin variant, but not AAV serotype, determines the opposite behavioral outcome of optogenetic stimulation of MeApd GABA neurons. Quantification of time spent attacking during 30 s immediately before-, or during light train stimulation (grey and blue bars, respectively), for all experimental conditions. The leftmost, and the second datasets (AAV8:ChETA and AAV2:ChR2 respectively) are re-plotted from Figures 1H and 3G. The third and the fourth datasets show the other two combinations of channelrhodopsin variants and AAV-serotypes: AAV2:ChETA (N=9), and AAV8:ChR2 (N=8 mice); example behavior traces are shown in Suppl. Figure 4. The two rightmost datasets represent the pooled AAV8 and AAV2 control groups, in which blue-light stimulation was ineffective (N = 11 and N = 7 mice, respectively). Note that irrespective of AAV serotype, optogenetic stimulation with ChETA resulted in *inhibition* of aggression, whereas optogenetic stimulation with ChR2 resulted in *stimulation* of aggression. See also Suppl. Figures 3, 4.

Taken together, the experiments in Figures 1 - 4 show that optogenetic stimulation of MeApd GABA neurons with ChETA versus ChR2 produce opposite behavioral outcomes: an *inhibition* of aggression with ChETA, versus a *stimulation* of aggression with ChR2 (Figure 4). This, obviously, presents a problem for the interpretation of these optogenetic stimulation experiments, because it cannot be assigned whether MeApd GABA neurons have a *stimulatory* - or else a *suppressing* role - in aggression control.

### ChR2 causes large plateau depolarizations, AP amplitude decrements and increased local inhibition

We next investigated the membrane potential changes driven by ChETA and ChR2 in MeApd GABA neurons, in order to correlate possible differences with the opposite behavioral outcomes of the two channelrhodopsin variants. For this, we expressed ChR2, or ChETA in a Cre-dependent manner using AAV2:EF1α:DIO:ChETA-eYFP and AAV2:EF1α:DIO:ChR2-eYFP (the same AAV2 vectors as used above for the behavioral experiments). In addition, we expressed cytosolic eYFP with a second, co-injected AAV2 vector (see Materials and Methods) to facilitate the targeting of transfected neuron in subsequent slice electrophysiology recordings, which were performed at near-physiological temperature (34-36°C).

We first recorded the responses of MeApd neurons to short trains of blue light (5 ms pulse length, 20 Hz, duration of trains 1s). Under voltage-clamp, we observed robust light-induced currents (Figure 5A, B; top) which had larger amplitudes in neurons expressing ChR2 - than ChETA (Figure 5C; p < 10^-4^; Mann-Whitney test), in keeping with previous work (Berndt et al., 2011; Mattis et al., 2012). The decay of the light-evoked currents was faster for ChETA than for ChR2 (p < 10^-4^ ; Mann-Whitney test; Figure 5D), as expected based on the engineering of the faster ChETA variant (Gunaydin et al., 2010). With repeated light stimuli, ChETA- and ChR2-expressing neurons fired APs reliably during the 1s trains (Figure 1A, B; middle). In ChR2-expressing neurons, however, a high plateau depolarization of 30.4 ± 1.9 mV was observed (analyzed over the first second of stimulation; n = 39 recordings; Figure 5E), significantly larger than in ChETA-expressing neurons (7.7 ± 0.9 mV; n = 25 cells; p < 10^-4^; two-tailed t-test; Figure 5E). One consequence of the plateau potential was a time-dependent decrease of the AP amplitudes from the first few stimuli onwards in ChR2-expressing neurons (Figure 1B, arrow), which was likely caused by voltage-dependent Na^+^-channel inactivation. An analysis of the time-course for AP amplitudes during the first 1s of light stimulation trains showed a significant effect of time in both the ChETA and ChR2 groups (p < 10^-4^ for ChETA and ChR2; one-way repeated measures ANOVA followed by a linear trend test), and the AP-amplitudes were significantly smaller in the ChETA group as compared to the ChR2 group (difference of ∼ 17 mV; p = 0.0096 for the channelrhodopsin variant effect; two-way repeated measures ANOVA followed by Holm-Šidák’s test for multiple comparisons; Figure 5I, left).

**Figure 5.**
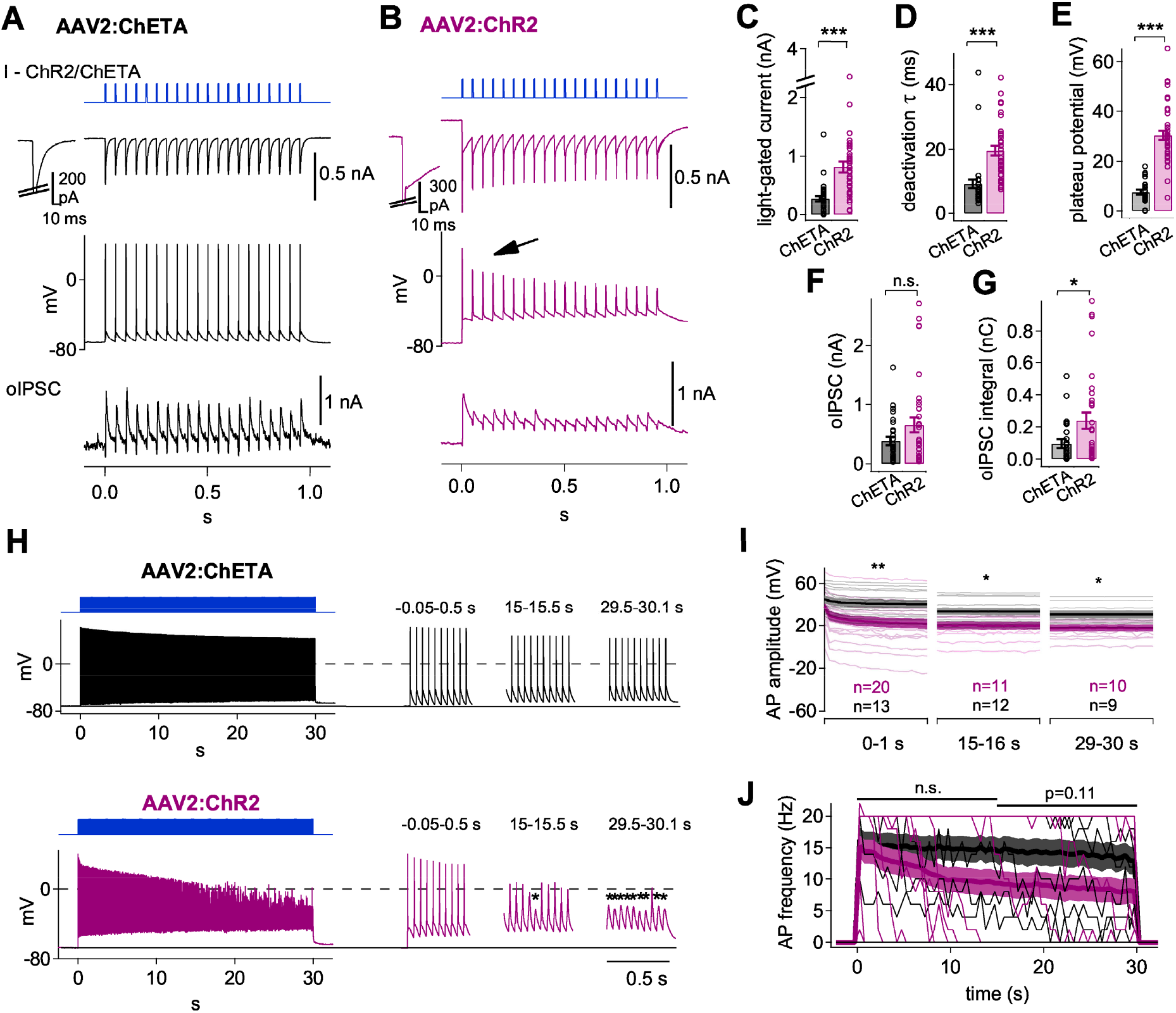
Optogenetic stimulation with ChR2 causes plateau depolarizations, AP decrements, and eventual failure of AP-triggering in MeApd GABA neurons. (A, B) Whole-cell recordings from example MeApd GABA neurons expressing either ChETA (A) or ChR2 (B). Cre-dependent expression was driven by the AAV2 vectors in VGAT^Cre^ mice (see Materials and Methods). Short (1s) trains of blue-light stimuli were applied (5 ms duration; 20 Hz repetition rate). Traces from top to bottom show: light-gated currents recorded at -70 mV holding potential (insets show the response to the first pulse; unclamped Na^+^ current components are clipped); current-clamp recording of membrane potential response (AP firing); optogenetically-evoked IPSCs (oIPSCs) recorded at 0 mV holding potential. (C - G) For each channelrhodopsin variant, we quantified the light-gated current amplitudes (C), the decay time constant of the light-gated current currents (τ; D), the plateau depolarizations (E), the first oIPSC amplitudes (F), and the integral of the oIPSCs (G). For statistical tests and statistical significance, see Results. (H) Example traces of AP-firing induced by in-vivo like trains (5ms, 20 Hz, 30s duration) at near-physiological temperature (35°C), in a MeApd GABA neurons expressing ChETA (top) or ChR2 (bottom). Insets to the right show traces during 0.5 s stretches at the start, middle, and end of the blue-light trains (corresponding times are indicated). Note the stronger decay of AP-amplitudes, and eventual AP-triggering failures in the ChR2- expressing neuron compared to the ChETA-expressing neuron. (I) Analysis of peak AP amplitudes at three times during the 30s light trains. Note a more pronounced time-dependent decrease of AP-amplitudes in ChR2-expressing neurons at the start of the train, and that the peak AP-amplitudes remain significantly smaller for ChR2 than for ChETA throughout the train. For statistical significance, see Results. (J) Plot of average AP-frequency attained by 20 Hz light pulse stimuli with ChETA (black traces) and ChR2 (violet traces), quantified for 500 ms bins. Shown are both individual traces, and the average ± s.e.m trace for each group. During the second half of the trains, the difference in attained AP-frequency (∼ 10.1 and 14.6 Hz for ChR2 and ChETA, respectively) did not reach statistical significance (p = 0.11). Error bars or color shadings are ± s.e.m. See also Suppl. Figure 5.

GABA neurons in the medial amygdala establish local inhibitory connections (Bian et al., 2008; Keshavarzi et al., 2014). We therefore next recorded optogenetically-evoked IPSCs (oIPSCs) at the reversal potential of the ChR2/ChETA currents (0 mV; Figure 5A, B; bottom), to compare the amount of optogenetically evoked mutual inhibition of MeApd GABA neurons with each channelrhodopsin variant. The first oIPSC amplitudes were not significantly different between the two groups (Figure 5F; p = 0.224). However, oIPSCs decayed more slowly after individual light pulses in preparations expressing ChR2, as compared to ChETA (Figure 5A, B, bottom). To quantify this effect, we analyzed the charge transported during the 1s-stimulation trains, which revealed a significantly, ∼ 2.5-fold larger inhibitory charge in the ChR2 group as compared to the ChETA group (Figure 5G; p=0.029; Mann-Whitney test; n = 33 and 25, respectively). Thus, a difference in optogenetically-stimulated local inhibition might contribute to the opposite behavioral outcomes between the two channelrhodopsin variants (see Discussion).

We next applied longer trains of blue light pulses, designed to mimic the *in-vivo* stimulation conditions (5 ms pulse width, 20 Hz, duration of 30s; Figure 5H). We again found a strong plateau depolarization in ChR2-expressing neurons, both during the first second as well as during the last 10 s, and this plateau depolarization was significantly smaller with ChETA (p < 10^-4^; two-way repeated measures ANOVA with Šidák’s multiple comparison test for 20 - 30 s; n = 17 and 24 recordings). Also, the AP amplitudes decreased in ChR2-expressing neurons to a stronger degree than in ChETA-expressing neurons, such that AP-amplitudes were smaller with ChR2 than with ChETA at various time points of the 30s-trains (Figure 5I; p=0.0096, p=0.011, and p=0.0113 in the three indicated time ranges for the channelrhodopsin factor in a two-way repeated measures ANOVA followed by Holm-Šidák’s multiple comparison test). Note that in the time range of 0-1s, AP amplitudes at every time point except t=0 were significantly lower in ChR2 expressing cells (p<0.05). With ChR2, we often observed that during the second half of the train, the peak of the APs dropped below 0 mV and in many instances blue light pulses failed to trigger APs (Figure 5H; stars). Correspondingly, during the second half of the trains, the average AP-frequency attained by the 20 Hz stimulus trains was only about 10 Hz in the ChR2-expressing neurons (Figure 5J). This was lower than in the ChETA-expressing neurons (∼ 15 Hz), but the difference did not reach statistical significance (p = 0.11; two-way repeated measures ANOVA; Figure 5J). This data shows that with long trains of blue-light stimuli as applied *in-vivo*, i) the plateau depolarization caused by ChR2 persists and becomes even larger (data not shown), ii) AP amplitudes were significantly smaller for stimulation with ChR2 than with ChETA, and iii) the AP-failure rate is higher for ChR2 than for ChETA during long trains, but the latter effect did not reach statistical significance.

We furthermore measured the intrinsic AP-firing properties in response to step current injections, and classified the AP-firing of the recorded MeApd GABA neurons to six types (Suppl. Figure 5). We found two types of MeApd GABA neurons with remarkably low rates of AP-firing even with high current injections (Suppl. Figure 5, AP-firing types 1 and 2; see also Lischinsky et al., 2017; Matos et al., 2020). In these neurons, optogenetic stimulation under ChR2 was often not able to trigger APs during the second half of trains (Suppl. Figure 5C). Nevertheless, the data set was too small to test for possible differences in AP- following, stratified for the six different AP-firing types of the MeApd GABA neurons.

## Discussion

The MeApd contains a majority of GABAergic neurons and has a crucial role in the control of social behaviors (Aleyasin et al., 2018; Haller, 2018; Hashikawa et al., 2016; Hong et al., 2014; Padilla et al., 2016; Unger et al., 2015). Previous work has suggested that optogenetic activation of these neurons, using the slow channelrhodopsin^H134R^ variant (ChR2), *stimulates* inter-male aggression (Hong et al., 2014). However, we found that optogenetic stimulation of MeApd GABA neurons, initially using the faster channelrhodopsin- 2^H134R,E123T^ variant (ChETA; Gunaydin et al., 2010) produced the opposite behavioral outcome, i.e. an *inhibition* of aggression. We therefore compared the optogenetic activation of MeApd GABA neurons with ChR2 and ChETA in behavioral experiments side-by-side, using two widely-used AAV serotypes to express the channelrhodopsins. These experiments showed unambiguously that with ChETA, aggression was *reduced* during the time of light stimulation (Figure 1), whereas with ChR2, an *increase* in aggression was observed (Figures 3, 4). To our knowledge, this is the first demonstration of how a kinetically different channelrhodopsin variants used to stimulate the same neuronal population *in-vivo* can lead to diametrically opposite behavioral outcomes.

An *ex-vivo* electrophysiological analysis of optogenetic stimulation of MeApd GABA neurons showed two main differences between the two channelrhodopsin variants: ChR2- expressing neurons showed a pronounced plateau depolarization of ∼ 30 mV, and, probably as a consequence, a decrement of the AP-amplitudes triggered optogenetically. Plateau depolarizations have been observed previously with ChR2, and are a consequence of the slow light-off kinetics of ChR2 (Berndt et al., 2011; Gunaydin et al., 2010; Herman et al., 2014; Mattis et al., 2012). It is safe to assume that the decrease in AP-amplitudes, and the partial failure of AP-triggering late during the trains, are caused by inactivation of voltage-gated Na^+^-channels (Hodgkin & Huxley, 1952), driven by the elevated plateau depolarization in ChR2- expressing neurons. Nevertheless, this difference in AP-triggering between ChR2 and ChETA, which was more apparent in the second half of the *in-vivo*-like trains and which did not reach statistical significance, is probably not the most important explanation for the different behavioral outcomes between the channelrhodopsin variants. Hong et al. (2014) found with shorter optogenetic trains of only 15 s length (20 ms pulse duration, 20 Hz), that optogenetic activation of MeApd GABA neurons under ChR2 produced an almost immediate, but lasting increase of attacks (Hong et al., 2014; their Figure 1). In our experiments with ChR2, we could reproduce the stimulatory effect of blue light trains on aggression; nevertheless, we did not observe a similarly fast increase in aggression (Figure 3D) as was observed by Hong et al. (2014) (their Figure 1). This difference might lie in the fact that Hong et al. used longer blue light pulses than we did (20 ms vs 5 ms, i.e. 40% vs 10% duty cycle, respectively). Increased pulse duration might have caused even stronger plateau depolarizations than we measured here, and potentially could lead to a partial light-induced depolarization block of certain MeApd GABA neuron sub- populations, as was demonstrated for different cell types in other brain regions (Herman et al., 2014). Taken together, these considerations suggest that a difference between ChR2 and ChETA in the first 5 - 10 s of light train stimulation might more likely account for the behavioral differences. We also hypothesize that the more efficient recruitment of local inhibition by ChR2 (Figure 5G) might be among the factors that causes the behaviorally different outcomes within the first 5-10s of blue light stimulation.

The MeApd consists of a majority (∼70%) of inhibitory neurons. Many of these neurons are long-range projection neurons to hypothalamic and other targets (Bian et al., 2008; Choi et al., 2005; Keshavarzi et al., 2014). Concomitantly, local inhibition described before for the MeApv (Keshavarzi et al., 2014) also likely occurs in the MeApd; our finding of optogenetically-evoked oIPSCs is consistent with local inhibitory circuits. Thus, while long-range inhibitory projection neurons to hypothalamic targets might drive an inhibition of aggression (see below), optogenetic activation under ChR2 leads to a stronger local inhibition than ChETA (Figure 5F, G). Thus, it is conceivable that this local opposing inhibitory effect reverses the behavioral outcome during *in-vivo* optogenetic activation with ChR2. Indeed, it is well-known that optogenetic activation of inhibitory neurons in cortical circuits causes an *inactivation* of the corresponding cortical area (Li et al., 2019; Sachidhanandam et al., 2013), because of the local nature of inhibition in the cortex. In analogy, optogenetic activation of VGAT^Cre^ positive neurons in the MeApd is expected to cause local inhibition in addition to the likely long-range inhibitory role of MeApd GABA neurons.

Hong et al. (2014) suggested a *stimulatory* role of MeApd GABA neurons in aggression control, which is now the predominant view in the literature (see reviews by Aleyasin et al., 2018; Chen and Hong, 2018; Hashikawa et al., 2016; Lischinsky and Lin, 2020). Nevertheless, Hong et al. did not identify the postsynaptic targets of inhibition by MeApd GABA neurons. Considering, however, that MeApd GABA neurons make a connection to hypothalamic nuclei including the VMH (ventromedial hypothalamus) and PMv (ventral pre-mammillary nucleus (Canteras et al., 1995; Choi et al., 2005; Lo et al., 2019), and further considering that optogenetic stimulation of specific populations of excitatory neurons in the VMH or PMv leads to a *stimulation* of aggression (Chen et al., 2020; Falkner et al., 2020; Lee et al., 2014; Stagkourakis et al., 2018; Yang et al., 2017), it might be expected that optogenetic activation of MeApd GABA neurons *inhibits* aggression, as we have observed with ChETA. Indeed, Hong et al. (2014) had to postulate a *disinhibition* of VMH glutamatergic neurons following activity of MeApd GABA neurons in order to explain their observations; nevertheless, such a disinhibitory pathway has not yet been identified. In a study published after Hong et al. (2014), NPY-expressing neurons in the MeA, which were to a large part GABA neurons, were found to increase aggression via the BNST (Padilla et al., 2016). This pathway via the BNST has been discussed as a possible *disinhibitory* route from the GABAergic MeApd to the VMH, and it was proposed to explain the earlier results by Hong et al. (2014) (see review by Chen and Hong, 2018). However, the correspondence of NPY+ neurons and MeApd GABA neurons remains to be shown, and the mechanisms by which the BNST controls aggression is at present not well defined, although an involvement of the BNST in aggression control is evident (Chen et al., 2020; Masugi-Tokita et al., 2016; Miller et al., 2019; Nordman et al., 2020). Yet another study showed that optogenetic activation of an excitatory projection from the dorsal raphe nucleus to the MeA inhibits aggression (Nordman & Li, 2020). This finding is consistent with our observation of an inhibition of aggression after optogenetic activation of MeApd GABA neurons. Taken together, the possible long-range inhibitory, or else - disinhibitory output pathways via which MeApd GABA neurons control aggression in either direction, should be elucidated in future studies.

Taken together, the paradoxical opposite behavioral outcomes of the optogenetic “sufficiency” experiment under ChR2 versus ChETA, does not allow us at present to conclude whether MeApd GABA neurons are involved in a *stimulatory* (Hong et al., 2014), or else in an *inhibitory* control of aggression. Our study calls for caution when choosing channelrhodopsin variants for *in-vivo* optogenetic studies with mechanistic conclusions, maybe especially so when studying inhibitory projection neurons. Thus, we suggest that the role of the MeApd GABA neurons in aggression control should be newly addressed. Such studies should include carefully controlled optogenetic experiments, as well as the identification of relevant long-range output targets of MeApd GABA neurons. In addition, the *in-vivo* activity of projection-defined MeApd GABA neurons should be recorded. Such experiments might be able to more conclusively establish the role of MeApd GABA system in the control of aggressive behavior.

## Author Contributions

A.B. and O.K. performed the experiments and analyzed the data; O.K., R.S., and A.B. conceived the study and wrote the manuscript.

## Acknowledgements

We would like to thank Dr. Dayu Lin and Dr. Julieta Lischinsky (New York University) for sharing unpublished results on ChR2-mediated optogenetic stimulation in the MeA, and Dr. Bernard Schneider (EPFL) for help with custom packaging of AAV vectors. Fluorescent microscopy was performed at the Bioimaging and Optics Core Facility (BIOP) at EPFL. This work was supported by the Swiss National Science Foundation, “NCCR Synapsy: The Synaptic bases of mental diseases”, project #28; to RS.

## Materials and Methods

### Laboratory mice

All procedures with laboratory mice (*mus musculus*) were authorized by the Service of Consumption and Veterinary Affairs (SCAV), Canton of Vaud, Switzerland (authorizations VD2885.0 and VD2885.1). We used the VGAT^Cre^ mouse line, which is a VGAT-internal ribosome entry site (IRES)-Cre line (STOCK Slc32a1^tm2(cre)Lowl^/J; RRID:IMSR_JAX:016962), first described in Vong et al. (2011). In some experiments (Figures 1, Suppl. Figures 4H, 4I), we used VGAT^Cre^ × tdTomato mice, which were the offspring of a cross between VGAT^Cre^ line and a Cre-dependent tdTomato reporter mice Ai9 (B6.Cg-Gt(ROSA)26Sor^tm9(CAG-tdTomato)Hze^/J; RRID:IMSR_JAX:007909), described in (Madisen et al., 2010). VGAT^Cre^ mice were back-crossed to the C57Bl6/J line (RRID:IMSR_JAX:000664) for 2-3 generations; similarly, the tdTomato line was maintained on C57Bl6/J background. Healthy adult, virgin male mice were used in a resident-intruder aggression tests. The animals were group-housed in cages with a maximum of five individuals at the specific pathogen free animal facility at EPFL under a 12/12 hr light/dark cycle, with food and water ad libitum. The animals were separated into single cages one day before surgery. BALB/cByJ male mice (RRID:IMSR_JAX:001026) were purchased from Charles River (Écully, France) and were group-housed until used as intruder mice in the resident-intruder test.

### Viral vectors

To target channelrhodopsin and/or fluorophore expression to MeApd GABA neurons for *in-vivo* and *ex-vivo* experiments, we used Cre-recombinase dependent (FLEX/DIO; Schnütgen et al., 2003) adeno-associated viral vectors (AAV) of the serotypes 2/8, 2/2 and 2/1 (referred to as AAV8, AAV2, AAV1; see Table 1). We used two modifications of the original wildtype ChR2 (Boyden et al., 2005; Nagel et al., 2003): a speed-optimized ChETA^H134R,E123T^ (Gunaydin et al., 2010), and an enhanced ChR2^H134R^ (Nagel et al., 2005) version. For production of the AAV8:hSyn:DIO:ChETA-eYFP and AAV8:hSyn:DIO:eGFP vectors used for our initial experiments (Figure 1), the DNA plasmids were respectively derived from the plasmids pAAV:EF1α:DIO:ChETA-eYFP (plasmid #26968; Addgene, Watertown, MA, USA) and pAAV:EF1α:DIO:eYFP (Addgene plasmid #27056) by custom cloning, and verified by sequencing. These two AAV8 vectors were packaged by the lab of Dr. Bernard Schneider (EPFL).

Because the previously used custom-made AAV2:EF1α:DIO:ChR2-2A-hrGFP vector (Hong et al., 2014) was not publically available, we purchased a vector driving the Cre-dependent expression of ChR2^H134R^ as close as possible to Hong et al. (2014): AAV2:EF1α:DIO:ChR2-eYFP (see Table 1). Indeed, with this vector, and with AAV8:EF1α:DIO:ChR2-eYFP, we were able to reproduce the previously reported stimulatory effect on aggression (Hong et al., 2014; see Figure 4). DNA sequence alignment has confirmed that all the channelrhodopsin-encoding vectors used in our study had the sequences downstream of promoters identical to plasmids pAAV:EF1α:DIO:ChETA-eYFP (Addgene #26968) and pAAV:EF1α:DIO:ChR2-eYFP (Addgene #20298), respectively, which were originally donated by the lab of Dr. Karl Deisseroth. Other viral vectors were purchased from various suppliers (see Table 1).

### Stereotactic surgery procedures

VGAT^Cre^ male mice (8-10 weeks old) in cohorts of 5-6 animals were randomly assigned to a test or to a control group. They were stereotaxically bilaterally injected in the MeApd with an AAV vector encoding for a channelrhodopsin- fluorophore fusion construct (test group) or fluorophore only (control group). All procedures were identical between the control - and the test groups. Stereotaxic surgeries were made using a model 942 small animal digital stereotaxic instrument (David Kopf Instruments, Tujunga, CA, USA) under continuous anesthestesia (1-1.5% isoflurane in O_2_) and pre- operative local analgesia (a mix of bupivacaine and lidocaine subcutaneously). For additional analgesia, paracetamol was provided in the drinking water (1 mg/ml) starting one day before, and during five days post-surgery. For *in-vivo* optogenetic stimulation experiments (Figures 1, 3, 4; Suppl. Figures 1, 4), 200 nl of virus suspension was injected on each brain side into the MeApd with a pulled glass capillary at the following coordinates (in mm from bregma skull surface): ML ±2.3, AP -1.7, DV -5.3. Optic fiber implants, custom made of 200 μm core / 0.39 NA optic fiber (FT200UMT; Thorlabs Inc, Newton, NJ, USA; implant transmission >70% at 473 nm) as previously described (Sparta et al., 2012; Tang et al., 2020), were vertically advanced to 500 µm above the injection sites and secured to the skull with a dental cement cup (Jet denture repair package; Cat#1234FIB; Lang Dental, Wheeling, IL, USA), and an anchoring screw (Cat#AMS90/1B-100; Antrin Miniature Specialties, Fallbrook, CA, USA). For *ex-vivo* patch-clamp recordings (Figure 5; Suppl. Figure 5), the injected suspension (200 nl) consisted of a 1:1 mix of two viral vectors as indicated, and no optic fibers were implanted. For histological analysis of AAV2 and AAV8 vector tropism (Suppl. Figure 3), 100 nl of a 1:1 mix of indicated viral vectors was unilaterally injected into the left MeApd.

### Behavioral testing

Following the surgery, the mice were housed individually. 4-5 weeks after surgery, a resident-intruder test was performed under optogenetic stimulation. The testing was performed at the beginning of the dark cycle in the home cages of the operated mice under minimal ambient light levels. No change of bedding was made for a week before testing; enrichment materials (nesting material, plastic shelter and cardboard tunnel) were removed for the experiment. There was no pre-selection of animals based on their basal aggression or hierarchy rank. Prior to the resident-intruder test, the operated VGAT^Cre^ resident mice were habituated to handling (being held in experimenter’s hands, constraining and patch cord attachment). Furthermore, the mice were habituated to the head tethering imposed by the optic patch cords while exploring the home cage, during 4-5 daily sessions 15 min long each. The resident-intruder experiment consisted of a 5 min baseline when the resident animal was exploring its home cage alone, followed by introducing an unfamiliar BALB/cByJ male intruder (8-9 weeks old) for the next ∼25 min. Each mouse underwent two sessions of resident-intruder testing on subsequent days using different intruders; each intruder animal was used maximally twice. The behavior was continuously recorded with two synchronized CMOS cameras (acA1300-60gc; Basler AG, Ahrensburg, Germany) providing top- and side views, equipped with infrared long-pass filters and under infrared light illumination, at 30 fps under control of EthoVision XT 13 software (Noldus Information Technologies, Wageningen, The Netherlands). The same software recorded a TTL signal from a Master-8 stimulator (A.M.P.I., Jerusalem, Israel) representing the gate for the trains of blue light pulses. The blue light pulses were produced by a 473 nm diode pumped solid state laser (MBL-FN-473-150mW, Changchun New Industries Optoelectronics, Changchun, China) triggered with TTL pulse trains from the Master-8 stimulator, and directed towards the optic fiber implants through 200 μm core / 0.22 NA optic patch cords (FC connector to 1.25 mm ferrule; Doric Lenses, Quebec, Canada) attached to a 1x2 intensity dividing fiber optic rotary joint (Doric Lenses). For each mouse, the laser power was adjusted to obtain 10 mW at the tip of the optic fiber implant; in some experiments we adjusted to other light power levels (Suppl. Figure 1). Light stimuli were 5 ms long, repeated at 20 Hz. We used a regular pattern of light trains (30-60 s long) interleaved with dark periods of fixed length (60 s), to avoid possible experimental bias. Such a bias could arise if the experimenter would trigger the light train manually during a short aggression bout, resulting in a trend of the light trains starting towards the end of the aggression bout.

### Histology

For the post-hoc validation of channelrhodopsin expression and optic fiber placement above the MeApd (Figure 2, Suppl. Figure 2), the animals were deeply anesthetized with intraperitoneal injection of pentobarbital, and transcardially perfused with 4% paraformaldehyde in PBS. After 24h post-fixation in 4% PFA and dehydration in 30% sucrose in PBS, 40 μm thick coronal brain sections were cut using a freezing microtome (Hyrax S30; Carl Zeiss, Oberkochen, Germany). For analysing the distribution of AAV2- and AAV8-infected GABA neurons in the MeApd, one VGAT^Cre^ mouse was injected with a mix of AAV2:EF1α:DIO:eYFP and AAV8:CAG:FLEX:tdTomato viruses. Two other mice were injected with AAV8:EF1α:FLEX:mCherry instead of the latter vector (Suppl. Figure 3). The animals were sacrificed 3 and 4 weeks after injection, respectively, and coronal sections of the MeA were prepared as described above. Immunohistochemistry was performed using 1:1000 diluted chicken anti-GFP (ab13970; Abcam, Cambridge, United Kingdom; RRID:AB_300798) and 1:500 rabbit anti-RFP antibodies (ab62341; Abcam; RRID:AB_945213) to stain against eYFP and tdTomato or mCherry, respectively. As the secondary antibodies, goat anti-chicken Alexa-488 (A11039; Thermo Fisher Scientific, Waltham, MA, USA; RRID:AB_2534096) and donkey anti-rabbit Alexa-568 antibodies (A10042; Thermo Fisher Scientific; RRID:AB_2534017) were used at 1:200 dilution.

In all cases, free-floating brain sections were mounted on glass slides and embedded in DAPI containing mounting medium (Fluoroshield with DAPI; F6057-20ML; Merck/Sigma-Aldrich, Darmstadt, Germany).

### Image acquisition

Brain sections for post-hoc validation of fiber placements (Figures 2, Suppl. Figure 2) were imaged with a slide scanning microscope (VS120-L100; Olympus, Tokyo, Japan) employing a 10x/0.4 NA objective. To analyze the distribution of AAV2- and AAV8-transduced neurons (Figure 5, Suppl. Figure 3), tile images of up to 3 focal planes were acquired using an inverted confocal microscope (LSM 700; Carl Zeiss) equipped with a 40x / 1.3 NA oil objective (Figures 5A, 5B, Suppl. Figures 3A-3C), or with a 20x / 0.8 NA air objective (Suppl. Figures 3D-3I) and 405, 488 and 555 nm laser lines for exciting DAPI, YFP and tdTomato/mCherry, respectively. Image acquisition was done at the Bioimaging and Optics Platform (BIOP), EPFL.

### Patch-clamp electrophysiology

A mouse at a time was sacrificed by decapitation after a brief anaesthesia with 3% isoflurane in O_2_ (according to the authorized protocol; see above), and 300 µm thick coronal brain slices containing the MeApd were made with a Leica VT1000S vibratome (Leica Microsystems, Wetzlar, Germany). The experiments were done 4-5 weeks following injection with the respective virus mix (Figure 5; see Table 1). Slicing was done in N-methyl-D-glutamine (NMDG) based solution containing (in mM): 110 NMDG, 2.5 KCl, 1.2 NaH_2_PO_4_, 25 NaHCO_3_, 20 HEPES, 25 D-glucose, 5 sodium ascorbate, 2 thiourea, 3 sodium pyruvate, 10 MgCl_2_, 0.5 CaCl_2_ (pH 7.3; Ting et al., 2014). The slices were then kept in a holding solution containing (in mM): 92 NaCl, 1.2 NaH_2_PO_4_, 30 NaHCO_3_, 20 HEPES, 25 D-glucose, 5 sodium ascorbate, 3 thiourea, 3 sodium pyruvate, 2 MgCl_2_, 2 CaCl_2_ (pH 7.3). Whole-cell patch clamp recordings were performed using an extracellular solution containing (in mM): 124 NaCl, 2.5 KCl, 1.2 NaH_2_PO_4_, 30 NaHCO_3_, 20 HEPES, 10 D-glucose, 5 sodium ascorbate, 2 thiourea, 3 sodium pyruvate, 2 MgCl_2_, 2 CaCl_2_ (pH 7.4, continuously bubbled with 95% O_2_ / 5% CO_2_). Recording pipettes were filled with a low-chloride solution, containing in (mM): 8 KCl, 145 K-gluconate, 10 HEPES, 3 Na-phosphocreatine, 4 Mg-ATP, 0.3 Na-GTP, 5 EGTA (pH 7.26, with KOH; 315 mosm). All recordings were done under an upright BX50WI microscope (Olympus) equipped with a 60x / 0.9 NA water-immersion objective (LUMPlanFl, Olympus), using an EPC-10 patch-clamp amplifier under control of PatchMaster software (HEKA Elektronik, Reutlingen, Germany). A near-physiological temperature control of the recording bath (34- 36°C) was done using a PM-1 heated platform, a SHM-6 inline solution heater and a TC- 344B controller (Warner Instruments, Holliston, MA, USA). The slices were illuminated with a Dodt IR gradient contrast and imaged using a CMOS camera (C11440-52U; Hamamatsu Photonics, Hamamatsu City, Japan) under control of MicroManager software (Edelstein et al., 2014). To visualize the eYFP or tdTomato/mCherry fluorescence, the fluorophores were excited using custom-fitted high-power LEDs (CREE XP-E2 Royal blue, 465 nm, and Green, 535 nm, respectively; Cree Inc, Durham, NC, USA) controlled with a BioLED driver (BLS-SA02-US; Mightex Systems, Toronto, Canada), and imaged using appropriate dichroic mirrors and emission filters. Short pulses (5 ms) of a blue LED at maximal power (∼80 mW/mm^2^ at the focal plane with 60x objective) were used for exciting ChETA or ChR2. oIPSCs were measured at 0 mV holding potential (Figure 5).

### Analysis of behavior data

The behavioral data was manually scored using the video annotation tool Anvil (https://www.anvil-software.org/; Kipp, 2001). During analysis, the scorer was blind to the timing of the optogenetic light train. During manual scoring, each video frame was exclusively classified as one of 23 elementary behaviors of the resident mouse: 1) attack; 2) tail rattle; 3) catching with paws; 4) proactive approach; 5) sniffing/touching the intruder’s body; 6) sniffing the body while following the intruder; 7) mouse/nose sniffing; 8) anogenital sniffing; 9) anogenital sniffing while following; 10) tail sniffing; 11) licking the intruder; 12) mounting the intruder; 13) being sniffed by intruder; 14) being followed; 15) being attacked; 16) being licked; 17) escaping from intruder; 18) grooming; 19) digging; 20) moving at a distance from the intruder; 21) resting at a distance; 22) rearing; 23) active avoidance. Display and averaging of the resulting behavioral traces was done using custom routines in IgorPro 7.08 (WaveMetrics, Lake Oswego, OR, USA). We combined the traces of elementary behaviors (see above) into the following six groups using a logical “OR”: “aggression” (behaviors #1-3), “following” (#6, 8-9), “social contact” (#4, 5, 7, 10, 11), “asocial” (#18, 19, 22, 23), “passive” (#13-17) and “resting/moving” (#20, 21). Note that mounting (#12) was not included in any of these groups because we have virtually never observed spontaneous nor light-evoked mounting in these experiments. The resulting six behavior traces were aligned to the onsets of the blue light trains and averaged across repeated trains for a given mouse (see e.g. Figures 1E, 1G). To quantify the effect of blue light exposure (e.g. Figure 1E), we compared the relative time of aggressive behavior during the first 30 s of the light train, to the last 30 s of the preceding dark period. The results of this analysis were averaged for each animal between two consequent days of resident-intruder testing, yielding one final data point per animal in the summary bar plot graphs (e.g. Figure 1H).

About one third of the behavioral experiments, performed towards the end of the study, was analyzed with an automatic procedure as described by Nilsson et al. (2020), using top-view videos and a DeepLabCut (Mathis et al., 2018; run on Google Colab platform) - SimBA behavior classification framework. The predictions of behaviors generated by SimBA (Nilsson et al., 2020) were analyzed in IgorPro the same way as the manually scored behaviors.

### Analysis of illuminated brain areas

The slide scanner images of brain sections for post-hoc validation of fiber placement were manually aligned into a serial stack using FIJI TrackEM2 (Cardona et al., 2010; Schindelin et al., 2012). A custom IgorPro macro was used for navigation through the stack and for fitting a model of the optic fiber into the fiber track visible on the images, in order to calculate the 3D location of an idealized light cone emanating from the fiber, thereby taking into account the fiber tilt. For computing a cone-shaped 3D light cone, we used the core diameter and numerical aperture of the fiber, and assumed a 500 μm limit for the light propagation in brain tissue (Aravanis et al., 2007). The contours of a brain atlas (Franklin & Paxinos, 2013) were overlaid onto the images using Adobe Illustrator (Adobe Inc, Mountain View, CA, USA). Finally, the areas of the relevant brain nuclei, their overlap with the extent of the channelrhodopsin expression, and the percentage overlap with the light cone were quantified in IgorPro. Before calculating the fractions, the respective areas were summed across the serial brain sections to approximate the fractions of volume (see Figures 2B, 2C, Suppl. Figure 2B-2C).

### Analysis of AAV vector tropism

For analysis of distribution of AAV2- and AAV8- transduced neurons (Suppl. Figure 3), non-overlapping ROIs were manually drawn using FIJI on confocal tile images around each neuron expressing any of the reporters (eYFP or tdTomato/mCherry), and the mean intensities and centroid coordinates of these ROIs were imported into IgorPro for analysis. The cells were classified as either expressing, or non-expressing the fluorescent reporter protein when the mean ROI fluorescence intensity exceeded a threshold value set at 150% and 200% of the mean background fluorescence in the whole MeApd for the red and the green channels, respectively. Based on this classification, binary maps of reporter (co-)expression by individual cells were generated (Suppl. Figure 3).

### Electrophysiological data analysis

Electrophysiological recording data were imported into IgorPro for analysis using custom routines. Peak amplitudes of light-gated channelrhodopsin currents and of oIPSCs (Figures 5C-5F) were measured at -70 mV and 0 mV holding potential, respectively, in response to the first light pulse in a train (Figures 5A, 5B; the second from top and the bottom panels, respectively). For some cells, the first light pulse evoked an unclampable Na^+^-current; in those cases, the value of the light-gated current was read out at the point of the falling slope change (if apparent; see Figure 5B insets), or as the peak of the next well-clamped channelrhodopsin-mediated current. The ChR2 or ChETA- evoked plateau potential during the 20 Hz / 5 ms light trains, was quantified as the average voltage offset over resting V_m_, after low-pass filtering <10 Hz to remove the action potential (AP) transients. Deactivation time constants of the two channelrhodopsin variants (see Results) were estimated by single-exponential fits to the decay of light-gated currents upon the last pulse in a train. Synaptic charge transfer (Figure 5G) was estimated by integrating the oIPSC trace over 1 s long train of light pulses (Figures 5A, 5B, bottom panels) after the baseline current subtraction; cells with the negative integral values due to negligible IPSC amplitudes were excluded. Analysis of the APs during the 30s trains of light pulses (Figures 5H-5J) was limited to cells expressing the direct current >50 pA, which lead to exclusion of n=2 and n=4 cells from ChR2 and ChETA groups, reducing the sample to n=25 and 18 recordings, respectively. For the absolute AP peak amplitude measurements during 30 s light trains (Figures 5H, 5I), fast voltage transients time-locked to the light pulses were first classified as APs only if exceeded 20 mV threshold relative to the passive membrane response level; the latter was estimated by filtering the voltage trace <10 Hz in a procedure identical to determination of the plateau potential. Only the cells that displayed no failures during the indicated time periods were admitted for analysis in Figure 5I; note that the cell numbers contributing to the average traces dropped faster for ChR2 than for ChETA, respectively, due to the failures in ChR2 expressing cells. For quantification of the number of successful APs generated during optogenetic trains (Figure 5J), APs were detected based on a threshold of 0 mV (see dashed line in Figure 5H) and considered as failure if the AP peak amplitude dropped below this threshold during the train.

Classification of the cells into intrinsic AP firing types (Suppl. Figure 5) was performed manually by an observer, based on the cell membrane voltage response to the step current injections (from -100 to +320 pA in 30 pA increments relative to the holding current; Suppl. Figure 5B). For this, the passive and active (AP firing dynamics) membrane properties were first analyzed with custom routines in IgorPro to extract information on the time course of AP frequency during the current step depolarizations, and on the maximal firing frequency as a function of injected current (Suppl. Figure 5B). Passive membrane properties such as the input resistance and membrane time constant were also documented, as well as the presence of a negative voltage sag (quantified as a “sag ratio”). The following definitions of the firing types were then used for classification (see Suppl. Figure 5B for examples). <u>Type 1</u>: a neuron responded with only a single AP to depolarizing current steps, then the AP firing ceased. <u>Type 2</u>: a neuron that slowly adapts to low spiking frequency at small injected currents, but which can only generate a short burst (a few APs) upon intermediate to large current steps. <u>Type 3</u>: a neuron generating a brief high-frequency burst of APs which then rapidly adapts to a low (<50 Hz) steady-state frequency. <u>Type 4</u>: a neuron showed no clear adaptation of the AP firing frequency, and in some cases – augmentation of instantaneous AP frequency showing a maximum between the 2^nd^ and the 3^rd^ APs. <u>Type 5</u>: a high-frequency spiking neuron slowly adapting to the high (≥ 50 Hz) AP frequency steady state at intermediate injected currents, and expressing no negative voltage sag. <u>Type 6</u>: AP firing dynamics as for the type 5 neurons, but expressing a negative voltage sag (a signature of I_h_ current, in some cases leading to post-hyperpolarization rebound AP). Note that at the largest amplitudes of tested depolarizing current steps, some neurons of the types 5 and 6 ceased AP firing after a short period of AP frequency adaptation (Suppl. Figure 5B).

### Statistical data analysis

No prior sample size calculation was performed. Resident-intruder experiments with optogenetic modulation were usually performed in small cohorts at a time of N = 3 and N = 3 mice for the test group (expressing ChETA or ChR2) and for the control group (expressing eYFP or eGFP); experiments were repeated with several cohorts for each condition. Statistical tests were performed using GraphPad Prism 9 (GraphPad Software, USA). A Shapiro-Wilk test for normality was made to determine if the data was normally distributed before choosing a parametric or non-parametric test type. In case of two-sample datasets, either a paired or two-sample t-test, or Wilcoxon or Mann-Whitney U tests were performed as indicated in the Results (Figures 1, 3, 4, 5C-5G, Suppl. Figures 1, 5). To compare the timecourses of AP amplitudes (Figure 5I; n=20 data points per cell in each time range), two-way repeated-measures ANOVA was performed for each time range to analyse effects of the channelrhodopsin variant and of the time, followed by Holm-Šidák’s multiple comparison test. For the analysis of AP following (Figure 5J), the average firing frequency was calculated in two 15s bins for each group, and a two-way repeated measures ANOVA analysis was performed followed by Šidák’s multiple comparison test. Significance levels were reported in the Results text and additionally indicated in the Figures by star symbols according to: p < 0.05, *; p < 0.01, **; p < 0.001, ***.

### Data availability

The datasets and custom code generated during this study will be made available for download at a public repository (https://zenodo.org), accessible with a DOI: https://doi.org/10.5281/zenodo.4311847

## Supplementary Figures

**Suppl. Figure 1.**
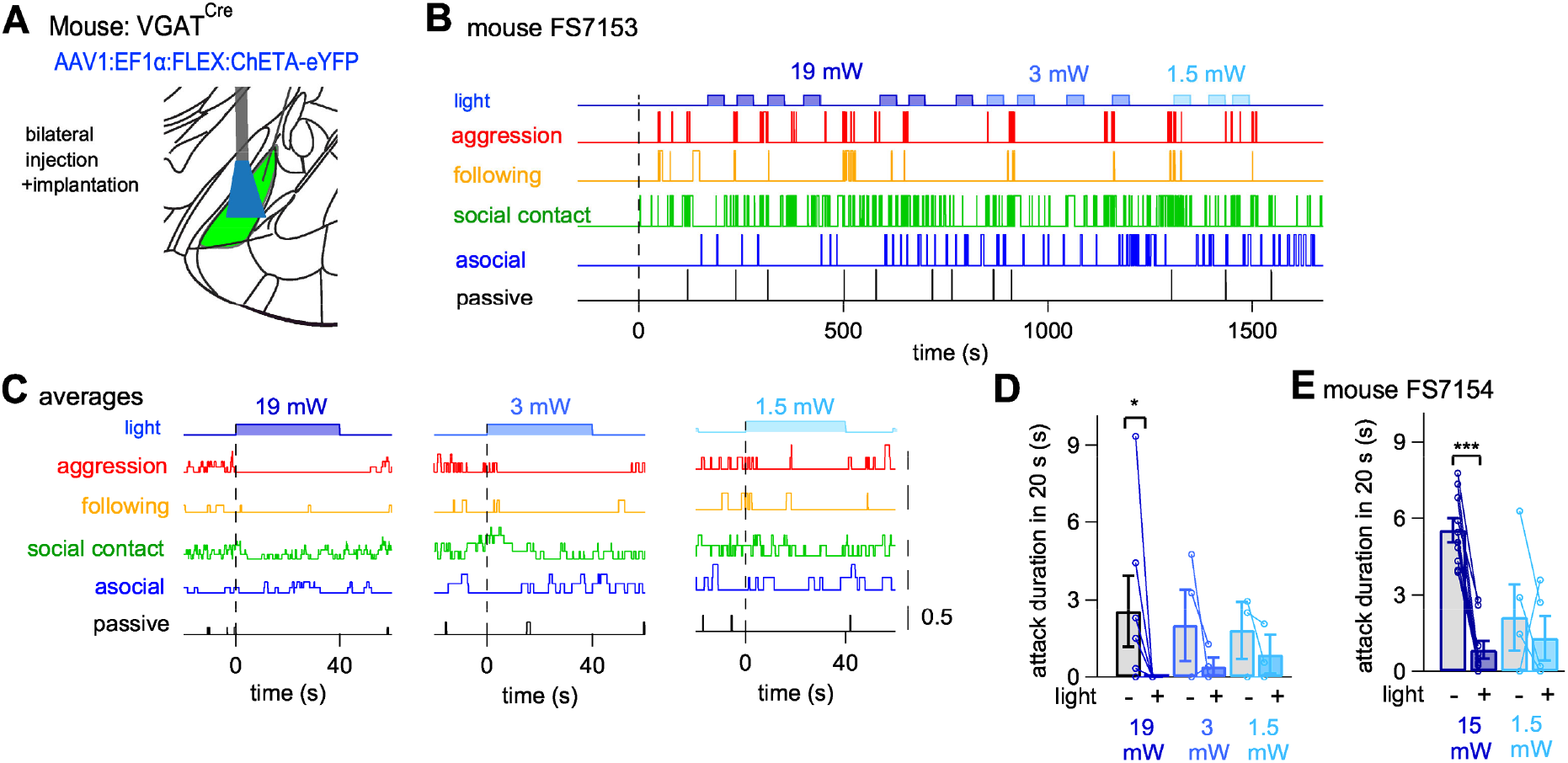
The inhibition of aggression by optogenetic activation of MeApd GABA neurons with ChETA depended on light power. Related to Figure 1. (A) Scheme of the approach for expression of ChETA and for optic fiber placement. In the experiments for this data set, an AAV1 vector was used to express ChETA. (B) Behavior traces during a resident-intruder test with a ChETA-expressing mouse. Trains of blue-light pulses (5 ms, 20 Hz, 40 s duration) were applied at varying intensities as indicated (19 mW, 3 mW and 1.5 mW; color-coded). (C) Average behavior traces aligned to the onsets of the light trains of different intensities (see B), as indicated. Note the clear interruption of ongoing aggression when using 19 mW light power and at 3 mW, whereas smaller light intensities of 1.5 mW did not have obvious effects. (D, E) Quantification of the time spent attacking during optogenetic stimulation with ChETA at different light intensities (D; same mouse as shown in B, C; for a second mouse see E). Note that significant differences (p = 0.03 and p = 0.001 in D and E, respectively; one-tailed Wilcoxon matched-pairs signed rank test) were observed at 19 mW and 15 mW, whereas behavioral changes were non-significant at lower light levels (3 and 1.5 mW). Error bars are ± s.e.m.

**Suppl. Figure 2.**
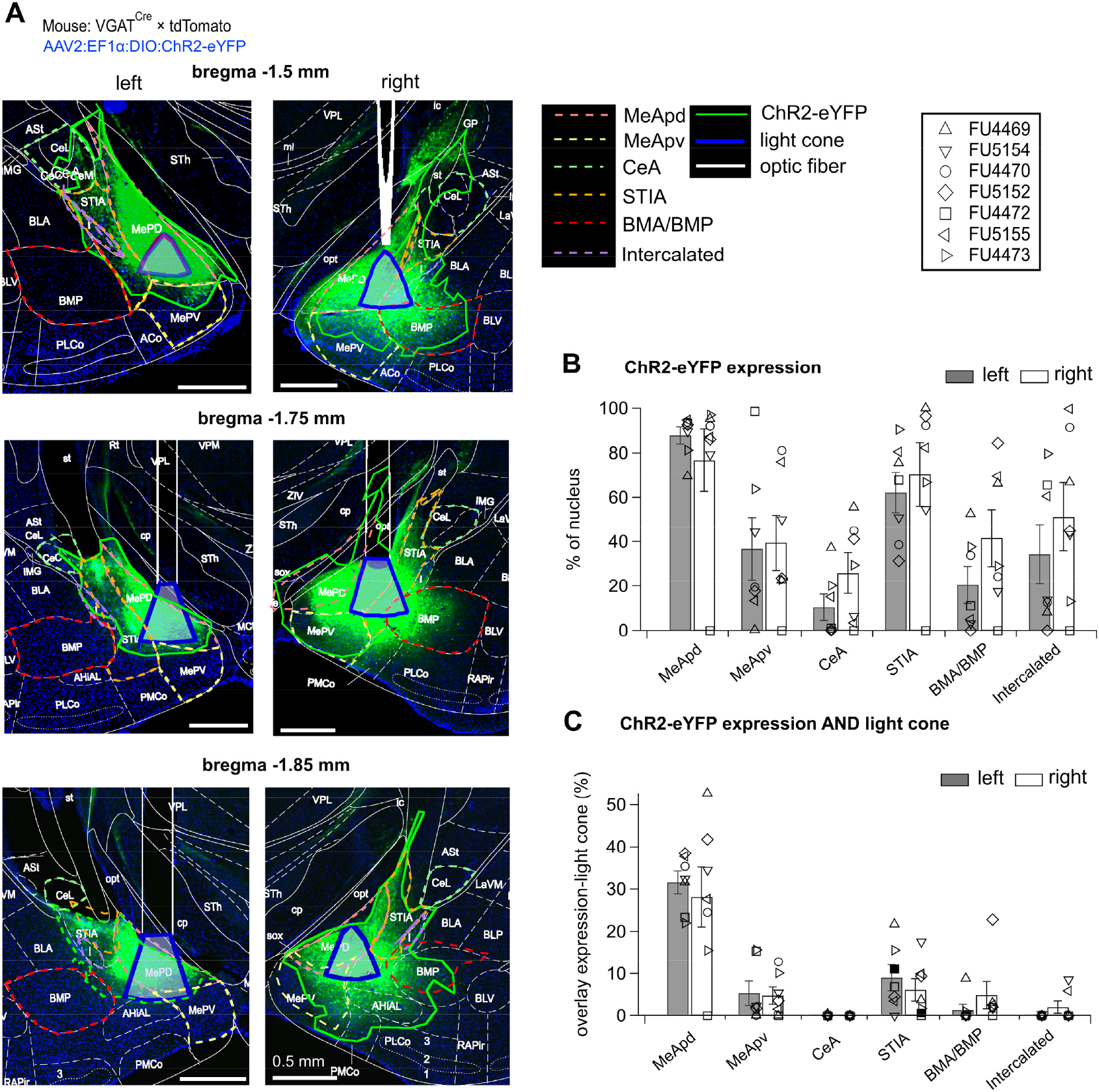
Histological analysis to identify the brain areas targeted by optogenetic stimulation with ChR2. Related to Figure 3. Here we performed a quantitative post-hoc histological analysis, analogous to that in Figure 2, in a cohort of N = 7 VGAT^Cre^ mice injected with AAV2:EF1α:DIO:ChR2-eYFP and subjected to a resident-intruder test with optogenetic stimulation of the MeApd GABA neurons (see Figure 3). The data of this histological analysis are presented in the same way as in Figure 2. Note that the results of the quantification (B, C) are very similar to those shown in Figures 2B, 2C for a cohort of ChETA expressing mice, and indicate that mainly the MeApd was targeted. We thus conclude that there were no gross differences in the targeting of the channelrhodopsin-variant expression, nor of the optic fiber placement, between optogenetic stimulation experiments using ChETA (Figure 1) and ChR2 (Figure 3). Error bars are ± s.e.m.

**Suppl. Figure 3.**
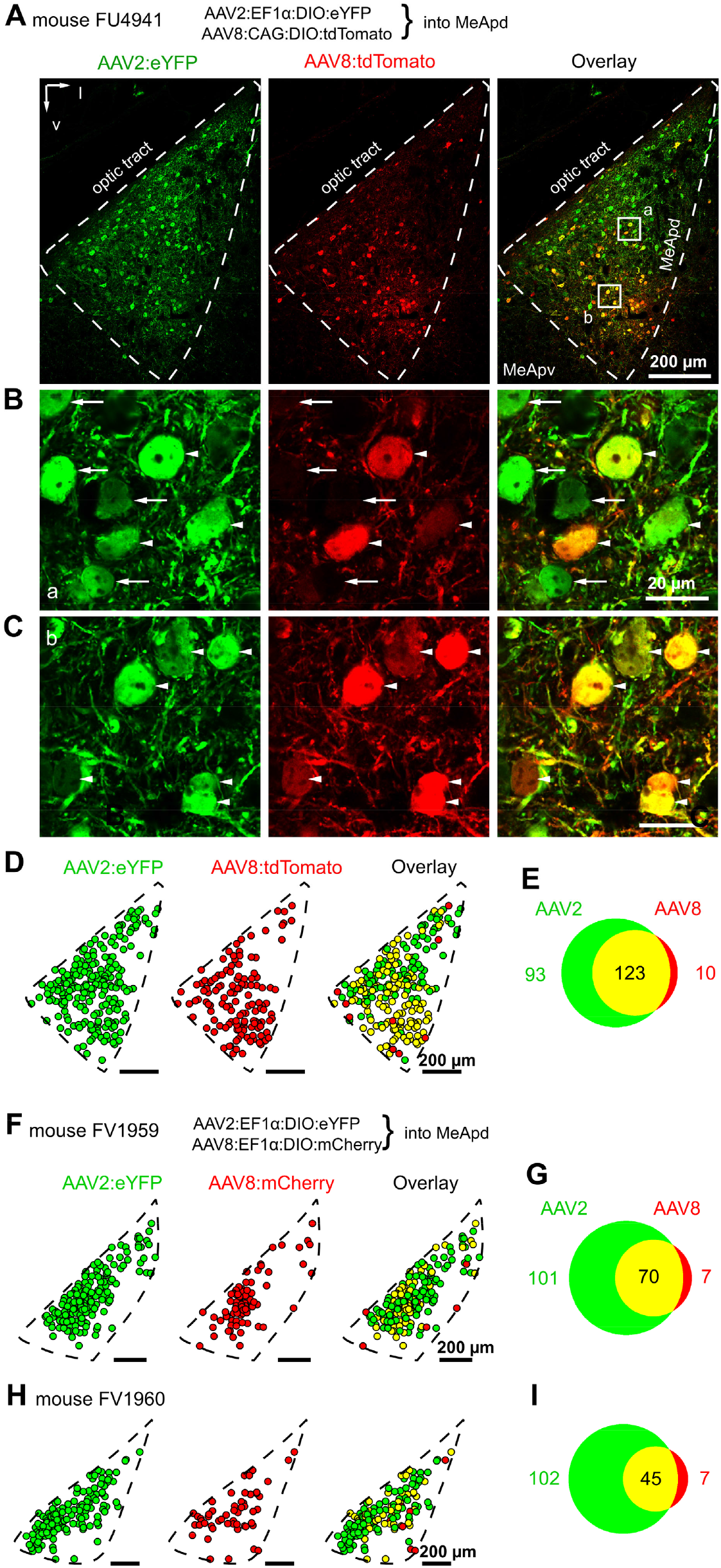
AAV8 vectors infect a sub-population of MeApd neurons transduced with AAV2 vectors. Related to Figure 4. (A) Overview confocal images of a coronal brain section containing the MeApd (white dashed outline), obtained from a VGAT^Cre^ mouse after injection with a mix of viral vectors AAV2:EF1α:DIO:eYFP (green channel; eYFP) and AAV8:CAG:DIO:tdTomato (red channel; tdTomato). Final titers of the injected vectors were 2.25·10^12^ and 3.25·10^12^ ml^-1^, respectively. (B, C) Zoomed-in images of the areas marked with the white rectangles in (A). *Arrows* point out example neurons expressing *only* eYFP from AAV2 (green channel); *arrowheads* point out neurons expressing tdTomato under an AAV8 serotype vector (red channel). The tdTomato-positive neurons often co-expressed eYFP (overlay, appearing yellow). (D) Binary maps showing the localization within the MeApd and identity of neurons in (A) that express either single marker (green or red), or both fluorescent proteins (yellow). (E) Venn diagram showing the proportions of neurons transduced by the AAV2 vector alone (green), by the AAV8 alone (red), or by both vectors (yellow). Same mouse as shown in (A-C); n = 2 sections were analyzed. (F - H) Binary maps and Venn diagrams as in (D, E), obtained from another two VGAT^Cre^ mice, four weeks after co-injection with AAV2:DIO:eYFP and AAV8:DIO:mCherry (n = 5 sections were analysed). Note the similarly of results obtained across N = 3 mice (D-I), which suggest that AAV8-transduced neurons are essentially a sub-population (∼ half) of the AAV2-transduced neurons. This finding is consistent with the behavioral findings, which did not reveal a fundamental difference in the behavioral outcome between the two AAV-serotypes, given that the same channelrhodopsin variant was used (Figure 4).

**Suppl. Figure 4.**
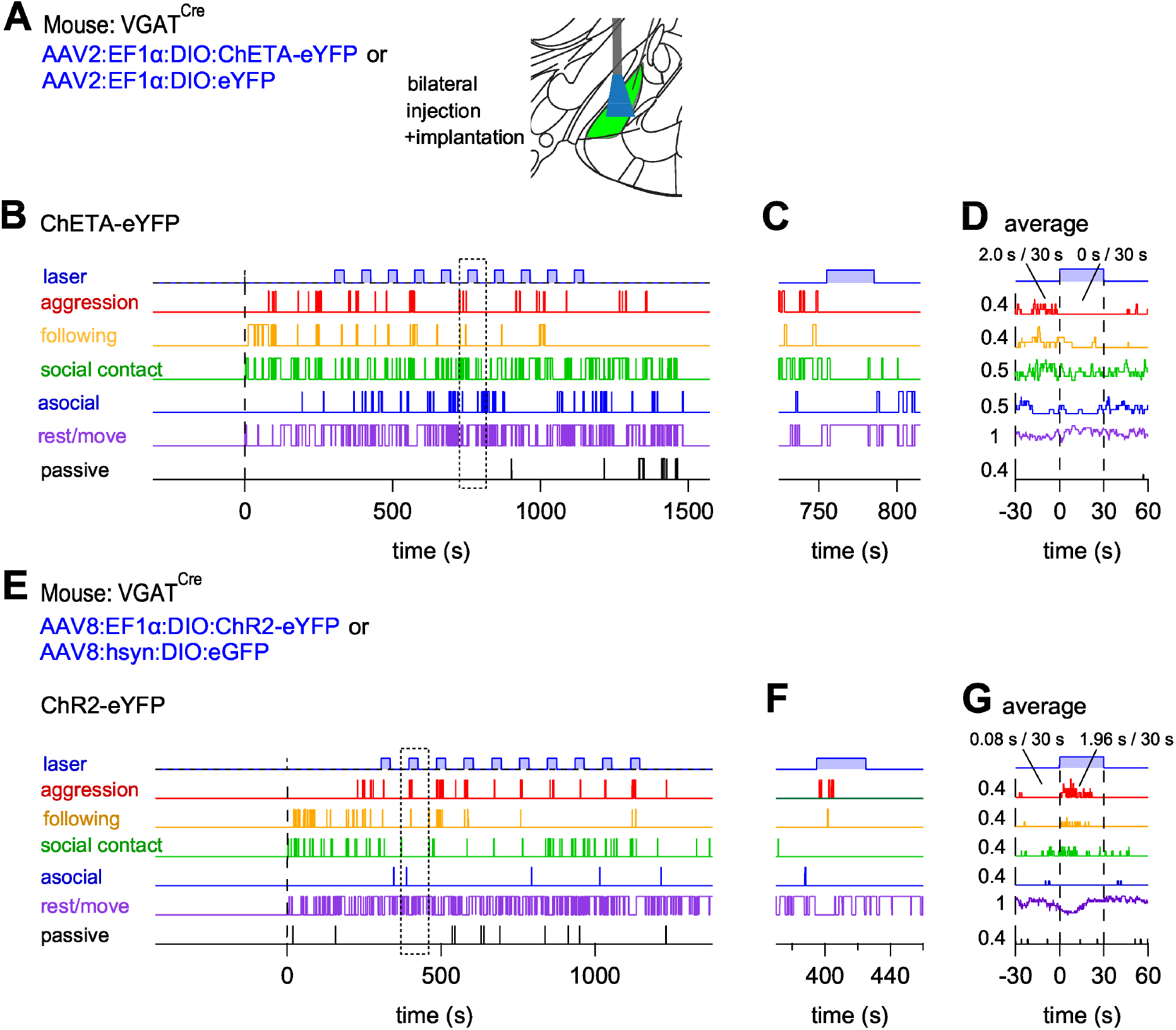
Example traces for optogenetic stimulation experiments under further combinations of AAV serotypes and channelrhodopsin variants. Related to Figure 4. Here, we show example experiments underlying the third and the fourth datasets in Figure 4 (AAV2:ChETA and AAV8:ChR2, respectively). (A) Schematic of the experimental approach to express ChETA, and for fiber placement. (B) Behavior traces during the resident-intruder test with a VGAT^Cre^ mouse expressing ChETA under an AAV2 serotype vector. (C) Example light episode marked with a dotted rectangle in (B). (D) Average behavioral traces from the experiment shown in (B), aligned to the onset of the light trains. Note the absence of aggressive behavior during the train of blue-light pulses. (E-G) Experimental data for a mouse expressing ChR2-eYFP under an AAV8 serotype vector. The expanded traces in (F) are taken from the area indicated by rectangle in (E). Note that aggression bouts preferentially occurred during the light trains (F), as reflected in the average aligned behavior traces (G).

**Suppl. Figure 5.**
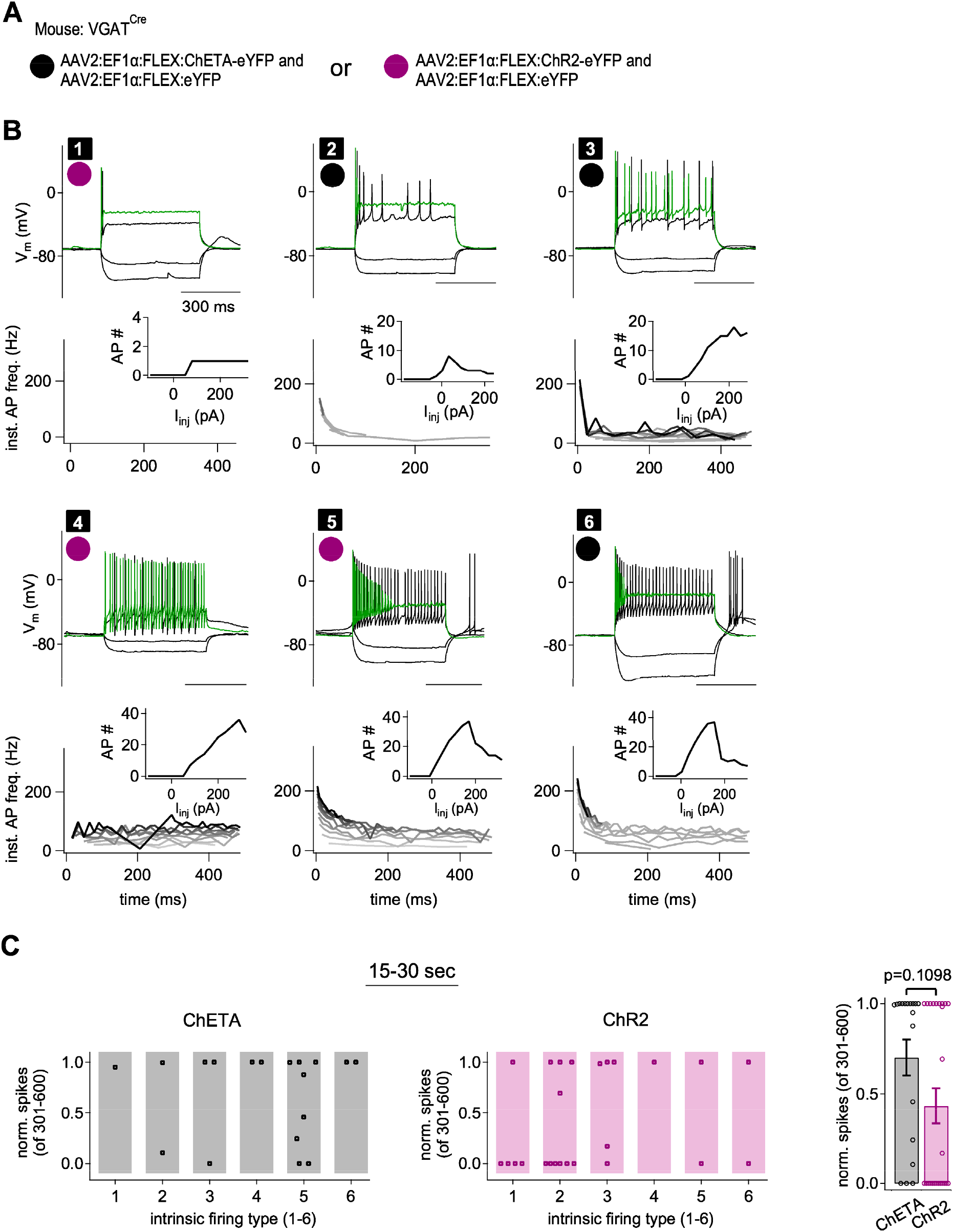
AP firing types identified amongst MeA GABA neurons, and distribution of channelrhodopsin variant - induced AP-firing across cell types. Related to Figure 5. (A) Experimental approach to express either ChETA or ChR2 in MeApd GABA neurons, co-expressing cytosolic eYFP from a co-injected second AAV2 vector for cell identification. (B) Examples of membrane potential (V_m_) responses of MeApd GABA neurons for each identified AP firing type (types designated 1 to 6; black squares). Responses to two hyperpolarizing (-100 and -40 pA) and two depolarizing current injections are shown (+110 and +290 pA; the latter in green). The expression of ChETA or ChR2 is indicated by black or magenta circle, respectively. The graphs at the bottom of each panel show the time-course of instantaneous AP frequency (darker traces show responses to larger depolarizing current steps), and AP-number as a function of the step current injection amplitude (insets). Materials and Methods describe the classification of the n = 6 AP-firing types in more detail. (C) Normalized number of APs during the second half of 30 s-long trains of blue light, segregated according to the AP-firing type of each neuron, both for MeApd GABA neurons expressing ChETA (left) and ChR2 (middle panel). Unfortunately, the number of recordings (n = 18 and 25 for ChETA and ChR2 in total) is too small to arrive at statistically meaningful difference in AP-following between the channelrhodopsins variants *and* across the n = 6 different AP-firing types. The quantification on the right shows the same data as on the left, collapsed over all AP-firing type cells in each group (p = 0.1098; Mann-Whitney test; see also Figure 5J).

